# A Tree of Human Gut Bacterial Species and its Applications to Metagenomics and Metaproteomics Data Analysis

**DOI:** 10.1101/2020.09.24.311720

**Authors:** Moses Stamboulian, Thomas G. Doak, Yuzhen Ye

**Author notes:** Corresponding author: Yuzhen Ye.

## Abstract

**Background:** Recent advances in genome and metagenome sequencing have dramatically enriched the collection of genomes of bacterial species related to human health and diseases. In metagenomic studies phylogenetic trees are commonly used to depict, describe, and compare the bacterial members of the community under study. The most accurate tree-building algorithms now use large sets of marker genes taken from across genomes. However, many of the current bacterial genomes were assembled from metagenomic datasets (i.e., metagenome assembled genomes, MAGs), and often contain missing information. It is therefore important to study how well the phylogeny approach performs on such genomes. Further, phylogeny methods are not perfect and it is important to know how reliable an inferred tree is.

**Results:** Here we examined the impact of incompleteness of the genomes on the tree reconstruction, and we showed that phylogeny approaches including RAxML (which handles missing data explicitly) and FastTree generally performed well on simulated collection of 400 genomes with missing information. As RAxML is computationally prohibitive for the much larger collections of gut genomes, we chose FastTree to build a unified tree of human-gut associated bacterial species (referred to as gut tree), including more than 3000 genomes, most of which are incomplete. We developed two downstream applications of the gut tree: peptide-centric analysis of metaproteomics datasets; and taxonomic characterization of metagenomic sequences. In both applications, the gut tree provided the basis for quantification of species composition at various taxonomic resolutions.

**Conclusions:** The gut tree presented in this study provides a useful framework for taxonomic profiling of human gut microbiome. Including MAGs in the tree provides more comprehensive representation of microbial species diversity associated with human gut, important for studying the taxonomic composition of gut microbiome.

**Availability and Implementation:** The tree construction pipeline and downstream applications of the gut tree are freely available at https://github.com/mgtools/guttree.

## 1 Background

Phylogenetic trees are by far the most widely accepted approach to represent organisms at various times throughout history and at various taxonomic levels, by illustrating the natural hierarchical relationships between different clades. Traditional methods in identifying and distinguishing between species often relied on morphological and other obvious behavioral characteristics in their classifications^1^. While such methods allowed for classification of many macroscopic organisms, it failed when applied to the microscopic world, especially bacteria and archaea^2^. With the rapid advancement of sequencing techniques now allowing for whole genome sequencing and reconstruction^3^, more systematic methods in (re)defining phylogenetic trees, based on molecular information, emerged. High-throughput Next Generation Sequencing methods^4^ coupled with Metagenomics made it possible to perform culture free genome sequencing, which removed biases such as sequencing only culturable species or reference genomes, further revealing the diversity of bacterial species^5^. These data sets allowed development of bioinformatics tools that computationally first assembled these (usually short) sequence reads^6,7^, binned them to complete or near complete genomes, and then assessed their quality^8–13^. All this has allowed for a rapid expansion in the number of unannotated genomes, many previously unknown, and associated with different environments, human body sites and geographic locations^14–17^.

Taking advantage of these developments and the expansion in complete and near complete genome databases, we focus our efforts on collecting a wide range of human gut bacterial genomes, in an attempt to characterize and catalogue the proliferation of different species across different individuals. Microbiomes, especially human gut bacteria, have received growing attention over the past decade. Numerous studies have been published focusing on human gut flora, reporting its important role in both human health and disease, most findings claiming its relationship to some sort of dysbiosis in the composition and functionality of gut microbes ^18^. Irritable Bowel Syndrome (IBS), a disorder characterized by abdominal pain, bloating, diarrhea or constipation, is thought to be caused by multiple factors, but variation in the composition of gut microbiota has been shown to be strongly associated with this syndrome as well ^19^. Inflammatory Bowel Diseases (IBD)—the most prominent ones being Crohn’s disease and ulcerative colitis—are characterized by inflammation of different parts of the intestines. It has been observed that alterations in the abundance and diversity levels of the Firmicutes phylum in the gut of some individuals contribute to IBD. The Fermicutes phyla in particular produce short chain fatty acids through fermentation of dietary fibers, and these acids have antiinflammatory properties^20^; patients diagnosed with IBD demonstrate a noticeable reduction in levels of Firmicutes. *Clostridium difficile* Infection (CDI) is another gut bacteria-associated disease, where hosts experience an increase in the gram-positive toxin producing bacteria *C. difficile* ^21^. It has been hypothesized that a dominant (healthy) gut microbiota protects their host from over-infestation of *C. difficile* species, through colonization-resistance mechanisms opposing it’s overgrowth^22^. These are just a few examples demonstrating the key role played by the human gut bacteria in regulating and maintaining the host’s health. Further work is now finding an interplay between gut microbiota and obesity and other metabolic syndromes^23^, allergic reactions^24^, and even with neurological disorders^25^. The importance of the human gut bacteria in health is indisputable, but understanding the mechanisms involved in maintaining a healthy homeostasis is just beginning.

Availability of genomic sequences dramatically broadens the diversity and resolution of the tree of life; genomics-based approaches are based on either a small number of marker loci, or use whole-genome information. Using more phylogenetic marker genes (e.g. a set of 16S ribosomal protein sequences from each organism)^1^ gives trees with higher-resolution than the 16S rRNA gene alone. Using this approach, Hug and colleagues derived a dramatically expanded tree of life, including over 1,000 uncultivated and often little known organisms^1^. But marker-gene based approaches require alignment of marker genes, which becomes difficult when the genomes are largely draft, each with its own incomplete set of marker genes. Alignment-free sequence analyses have been applied to problems ranging from whole-genome phylogeny to the classification of protein families, identification of horizontally transferred genes, and detection of recombined sequences^26^. Alignment-free approaches (such as kmer based methods) are more resilient to compositional biases, complex genetic rearrangements and large insertions/deletions for whole genome phylogenies^27–29^. CAFE is an efficient implementation of a fast kmer-based alignment-free method to calculate distances across genomes^29^. But alignment-free methods discard the great deal of information found in protein alignments. On the other hand, alignment based maximum likelihood or Bayesian approaches such as RAxML, IQ-TREE and Mr. Bayes can explicitly handle missing information^30,31^, but are usually computational expensive so cannot handle large alignments involving many sequences. It was shown that FastTree is more scalable, yet at the same time retains comparable performance to the slower programs ^30,31^. We also show that FastTree achieves a similar performance to RAxML on alignments with missing information, using simulated datasets with missing information (see results).

Recent efforts have focused in collecting gut bacterial genomes from shotgun sequenced metagenomic samples, representing different states of health and dysbiosis, and a collective 3,301 metagenome assembled genomes (MAGs) were recently^32^ and^33^. A tree of these species will provide valuable information on the biodiversity of gut bacteria, and will be practically useful for a spectrum of analytic methods that rely on the phylogenetic relationship of the species, including MetaGOmics^34^ and Unipept (https://unipept.ugent.be) ^35^. Here we report a comprehensive alignment-based phylogenetic tree of human gut-related bacterial species using FastTree over 120 ubiquitous bacterial marker genes, and explore its applications^36^. These marker genes have been shown to be present in *≥*90% of the bacterial genomes, as single copies in most cases^37^. The ubiquity and importance of these genes make them less likely to be subjects of horizontal gene transfers within prokaryotic species^38^. We show that alignment concatenation based approaches remain superior and result in better phylogenetic trees as compared to alignment-free methods, as tested on a subset of simulated species by deliberately removing part of their genomes. We discuss the challenges in constructing a tree representing evolutionary distant species, as well as challenges using nearly complete genomes with possible missing parts. We reflect on certain limitations and disagreements between our tree and the taxonomic assignment of the genomes based on a lowest common ancestor approach. Finally we explore applications of the constructed gut tree in the context of representing the most abundant and dominant species across different samples from metaproteomic and metagenomic perspectives and reflecting it on the tree. We believe that the gut tree we built and the companion tools we developed for using the tree will be beneficial as an initial attempt in profiling and performing comparative studies across different human subjects coming from diverse backgrounds and health conditions.

## 2 Methods

### 2.1 Gut bacterial genomes and metagenome assembled genomes (MAGs)

We collected over 3,000 genomes for human gut bacterial species from two recent studies^32,33^. These two studies report the largest genomic catalogue for human gut bacteria thus far. Bacterial genomes reported in^32^ were compiled from two sources: a total of 617 genomes obtained from the human microbiome project (HMP)^39^, and 737 whole genome-sequenced bacterial isolates, representing the Human Gastrointestinal Bacteria Culture Collection (HBC). These 737 binned bacterial genomes were assembled by culturing and purifying bacterial isolates of 20 fecal samples originating from different individuals^32^. On the other hand the bacterial genomes reported in^33^ were generated and classified from a total of 92,143 metagenome assembled genomes (MAGs), a total of 1,952 binned genomes were characterized as non-overlapping with bacterial genomes previously reported, i.e., don’t match with any isolate within UniportKB. These novel binned genomes were termed as Uncharaterised MetaGenome Species (UMGS). We were able to retrieve 612 out of 617 RefSeq sequences using the reported RefSeq IDs. Our final dataset for this study consists of 3,301 genomes and MAGs, inclduing 612 genomes from the RefSeq database, 737 whole genome-sequenced bacterial isolates from the HBC dataset and 1,952 UMGS genomes.

### 2.2 Gene prediction and marker gene assignment

For RefSeq genomes, we obtained their genes from the RefSeq database: a total of 1,907,611 genes were retrieved for all the RefSeq genomes included. For the rest (i.e., the MAGs), we applied FragGeneScan (FGS), with default settings, to predict protein coding genes from the contigs^40^: a total of 2,602,889 genes were predicted from the HBC genomes, and another 4,001,749 genes from the UMGS genomes.

We used a set of a set of 120 marker genes for phylogeny reconstruction. We extracted hidden markov models (HMM) for these 120 marker genes from Pfam^41^ and TIGRfam databases^42^ (the list of HMM models can be found here). We then applied hmmscan (in the hmmer3 package) to search against the HMM models, to predict marker genes in the genomes and MAGs, with an e-value of *e*^*−*10^ as the cutoff.

### 2.3 Data simulation and performance comparison of phylogeny approaches

To compare different phylogenetic inference methods (FastTree and RAxML) and clustering methods (alignment free based), we used a subset of 400 of the 612 gut-associated RefSeq genomes sharing 57 pfam domains (see here for a comprehensive list of shared Pfams). FastTree and RAxML approaches are alignment-based. HMMalign was used to construct sequence alignments for each of the 57 domains^43^. G-blocks was used for the final alignment concatenation and improvement^44^. Default settings for RAxML and FastTree were used to infer phylogenetic trees over this dataset. For the alignment-free approach CAFE, we used two distance measures, the Manhattan distance and D2S^29^. Calculating the Manhattan distance across genomes is computationally less demanding in comparison to the other distance measures available, and we employed it here mainly to optimize parameters, such as k-mer size, using whole genomes as opposed to protein coding genes or highly expressed genes. We chose k-mer size of 8, based on our experimenting, and it is consistent with the authors’ recommendation for computing the evolutionary distances between bacterial genomes^29^. It should be noted that whole genome sequences were used in the case of alignment free methods.

Simulation of missing data was performed by uniformly randomly selecting 100 of the 400 genomes (25%) from each of the individual multiple sequence alignment (for each Pfam) and removing those species from the alignment (by replacing their respective alignments with gaps). This was done to mimic the absence of some of marker genes in certain genomes due to either incompleteness of the MAGs, or due to gene loss events in certain lineages, simulating the effects of genomes with missing marker genes.

Given a phylogenetic tree, pairwise species evolutionary distances were calculated using the cophenetic function from the APE R package ^45^. We then use the correlation (both Pearson and Spearman correlations) between the pairwise species evolutionary distances to quantify the similarity between the different phylogenetic trees built from the same set of species. We didn’t use tree-based metric such as RobinsonFoulds metric for tree comparison, since it saturates rapidly so very similar trees can have large distance value^46^.

### 2.4 Constructing the tree of gut bacteria

All 3,301 genomes/bins were used to construct a comprehensive bacterial tree. From each genome or MAG the annotated and predicted protein sequences of the marker genes were extracted. A total of 73,285, 87,742 and 210,374 proteins from RefSeq genomes, HBC bins and UMGS bins were extracted respectively. The aligned regions between each protein sequence and HMM models were extracted. A total of 120 individual multiple sequence alignments were constructed between sequences extracted by each HMM model using hmmalign ^43^. Gaps were used in the alignments for genomes with missing marker genes. Individual alignments from the 120 domains were then concatenated and further refined by removing columns with more than 50% gaps, columns with a consensus of less than 25% and rows with more than 50% gaps. A phylogeny was inferred by constructing a species tree from the final concatenated multiple sequence alignments using FastTree under WAG + GAMMA models^36^. We used -pseudo, -spr 4, mlacc 2 and -slownni options while running FastTree to increase the accuracy of the tree inference^36^. The final tree was rooted using eight genomes belonging to the family *Saccharimonadaceae* as an outgroup. We chose this clade since they were the only representative genomes in the *Patescibacteria* phylum. Branch support values were also calculated by using 100 bootstrapped trees replicates. Bootstraps were calculated using the *seqboot* function from PHYLIP tools^47^.

### 2.5 Downstream applications of the gut tree for metagenomic and metaproteomic data analyses

We developed two downstream applications for tree-based metaproteomics and metagenomics analyses, providing quantification of the relative abundances of different bacterial species at different taxonomic resolutions based on the metagenomic or metaproteomic data. For metagenomic data analysis, our pipeline applies Bowtie2 to map sequencing reads to the collection of gut genomes. Based on the reads mapping results, we implemented two methods for genome abundance quantification. The first method is the Lowest Common Ancestor (LCA) approach^48,49^ which assigns reads to the nodes in the tree such that if a read maps to multiple genomes, it is assigned to their lowest common ancestor. We also implemented an alternative quantification approach using all mapped reads (uniquely and multi-mapped), such that multi-mapped reads are assigned to genomes proportionally according to their abundances computed using only uniquely-mapped reads (called *multi-mapped approach*)^50^. We tested our pipeline using a publicly available gut metagenomic dataset^51^ and compared our results to those from MetaPhlAn2^52^ and Kraken2^48^.

For metaproteomic data analysis, given identified peptides, our pipeline first maps the peptides to the proteomes of the gut genomes, and then applies the LCA approach to infer the relative species abundances at various taxonomic levels (the multi-mapped approach isn’t applicable due to the low throughput of metaprotomics approaches). We applied our pipeline to analyze sample peptides identified from a human gut microbiome available on Unipept’s website, and compared our results to the results from Unipept (version 4.3) ^53^.

### 2.6 Availability of the tree and companion tools

The gut tree and companion tools are available at https://github.com/mgtools/guttree. Data required for the applications of our tools, including the genome sequences (for reads mapping and following taxonomic assignment and quantification from metagenomic data) and protein sequences (for peptide-centric metaproteomics analysis) are available for download at http://omics.informatics.indiana.edu/guttree/.

## 3 Results

We first report the comparison of tree reconstruction methods using simulated datasets with missing information. We then report the tree of human gut bacteria built using the selected approach. Finally we demonstrate applications of the tree in two downstream analyses.

### 3.1 Testing different phylogeny approaches on simulated datasets with missing information

To evaluate the efficacy of different phylogeny approaches for tree reconstruction using incomplete genome information, we used 400 RefSeq genomes as described in the methods section. We simulated datasets from these 400 RefSeq genomes with missing data by removing genes from them. On average, the gut genomes each miss 7% of their marker genes (see Supplementary Figure 1, but some may be more incomplete. We therefore chose to randomly remove a significant fraction, 25% of the sequences from the MSA to simulate genomes with missing information. FastTree, RaxML and CAFE (alignment-free approach) were applied to build trees of these genomes with and without missing information using different approaches. For CAFE, two distance measures, Manhattan and D2S, were tested.

Results show that FastTree and RAxML gave similar trees, and they outperformed the alignment-free approaches. Figure 1A and 1B show comparisons between FastTree and RAxML, on complete genomes, and missing genomes, respectively. Species pairwise phylogenetic distances were highly correlated between the two approaches over the complete genomes (Pearson correlation of 0.983 and Spearman correlation of 0.942). Surprisingly, an almost perfect correlation was achieved when both of these algorithms on the dataset with simulated missing markers (Pearson correlation of 0.996 and Spearman correlation of 0.989). For alignment-free approaches, the D2S measures outperformed the Manhattan distance, which is consistent with previously reported results^29^. Pearson and Spearman rank correlations of 0.569 and 0.219 respectively were observed between the D2S k-mer based distances and the pairwise distances between species computed from FastTree phylogenetic tree (see Figure 1C); by contrast, the correlations marginally dropped to 0.513 and 0.250 respectively when Manhattan distance was used to compute the k-mer based distances (see Supplementary Figure 2). Similar trends were observed when we repeated the experiments by comparing the alignment-free approaches to the “ground truth” constructed using RAxML instead. Detailed results of this experiment are summarized in Supplementary Figure 3. On the other hand, when comparing correlation coefficients between the distance matrices calculated from FastTree trees with and without missing data, we obtained a near perfect Pearson’s correlation of 0.982 and a significantly higher Spearman rank correlation of 0.940, as shown in Figure 1D.

**Figure 1:**
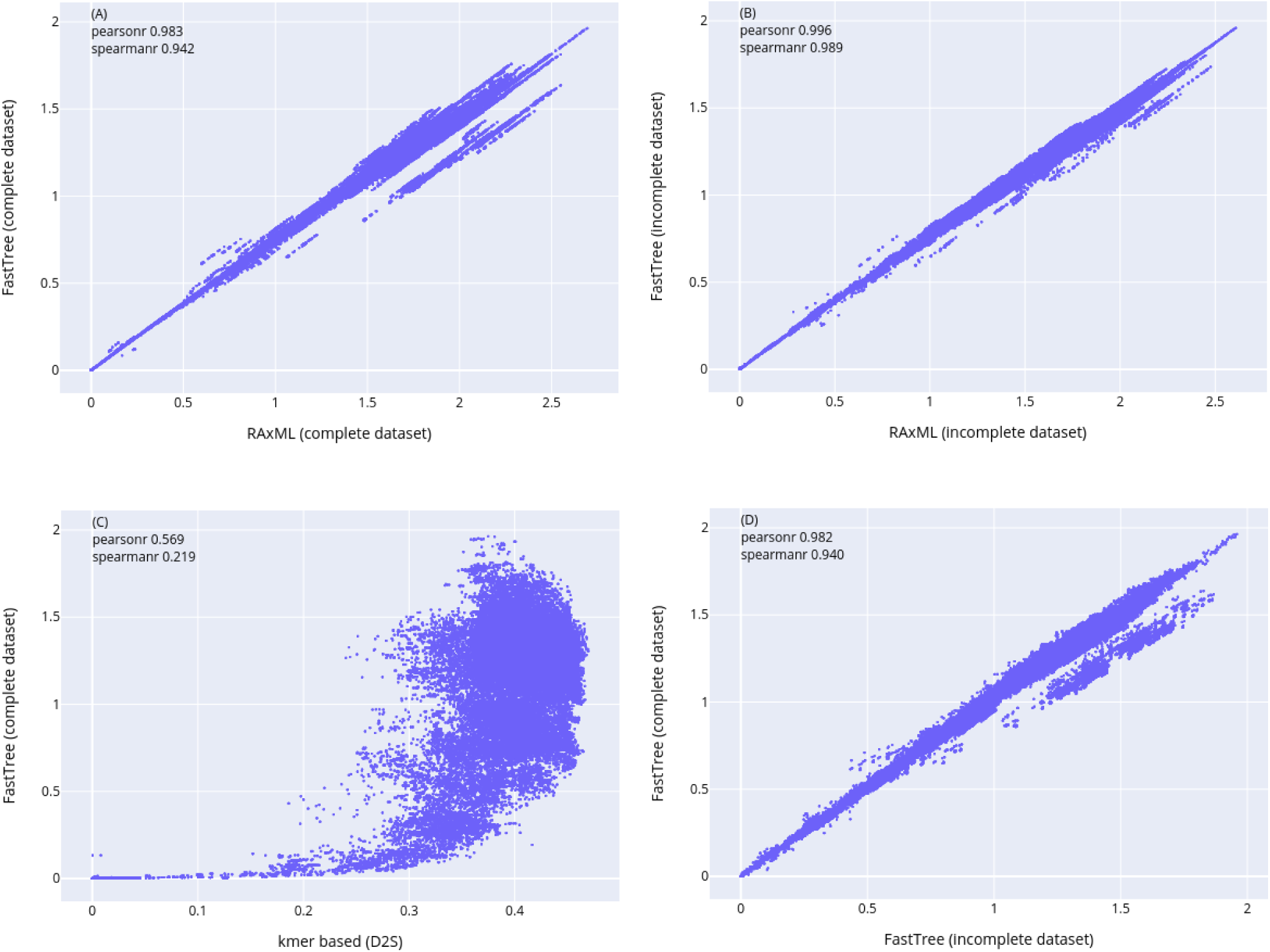
Correlations of pairwise species distances derived by different approaches. (A) RAxML and FastTree on 400 RefSeq genomes. (B) RAxML and FastTree on simulated genomes with missing marker genes. (C) Alignment free method using D2S distance and FastTree using complete genomes. (D) FastTree trees on complete genomes vs genomes with missing marker genes.

### 3.2 Tree of human gut bacteria

Based on our simulation results (above) and also the computational efficacy of the methods, we chose FastTree as the method to reconstruct a comprehensive tree of gut bacteria—most of which have incomplete genomes—using all the available sequences with significant hits to the Pfam and TIGRfam domains for gene alignments from each genome: this resulted in the largest and most representative human gut-associated bacterial tree by far, to the best of our knowledge. All but eight genomes participated in the final tree construction: we removed eight MAGs from participating in the final tree construction, due to their poorly aligned regions, which did not meet the multiple sequence alignment criteria used to construct this tree (see Methods for more details). Distributions of the completeness and the contamination levels of the bins participating in constructing our gut-tree are summarized in Supplementary Figure 4.

Out of the 120 Pfam/TIGRfam marker profiles used, astonishingly, only three marker profiles were shared across all the genomes, which were leucine tRNA ligase, translation initiation factor and GTPase domains. We categorized these 120 protein functional domains into three general groups: *ribosomal proteins, enzymes* and *transcription related proteins* with quantities 30, 61 and 29 domains respectively. On average, each genome/MAG had about 26 different ribosomal protein domains, 58 enzyme domains and 27 transcription related proteins. These distributions fluctuate greatly between the three different sources of genomes. Individual statistics for NCBI, HBC and UMGS genomes are summarized in Figure 2. While the distributions of these categories across the three genome sources loosely follow similar trends, with small fluctuations, it varies significantly when comparing them to those of the UMGS genomes: the number of the different protein domains found in UMGS genomes varies more and on average is lower than that of NCBI and HBC genomes, further suggesting that the UMGS genomes are more incomplete than the other sources. On average each genome contains over 93% of these domains (a complete summary for the number of genomes containing each of the pfams can be found in Supplementary Figure 1). After removing poorly aligned columns and rows with the above mentioned criteria, a multiple sequence alignment of 3,293 rows (species) and 33,182 columns was used to infer a phylogeny using FastTree. The resulting tree is summarized in Figure 3.

**Figure 2:**
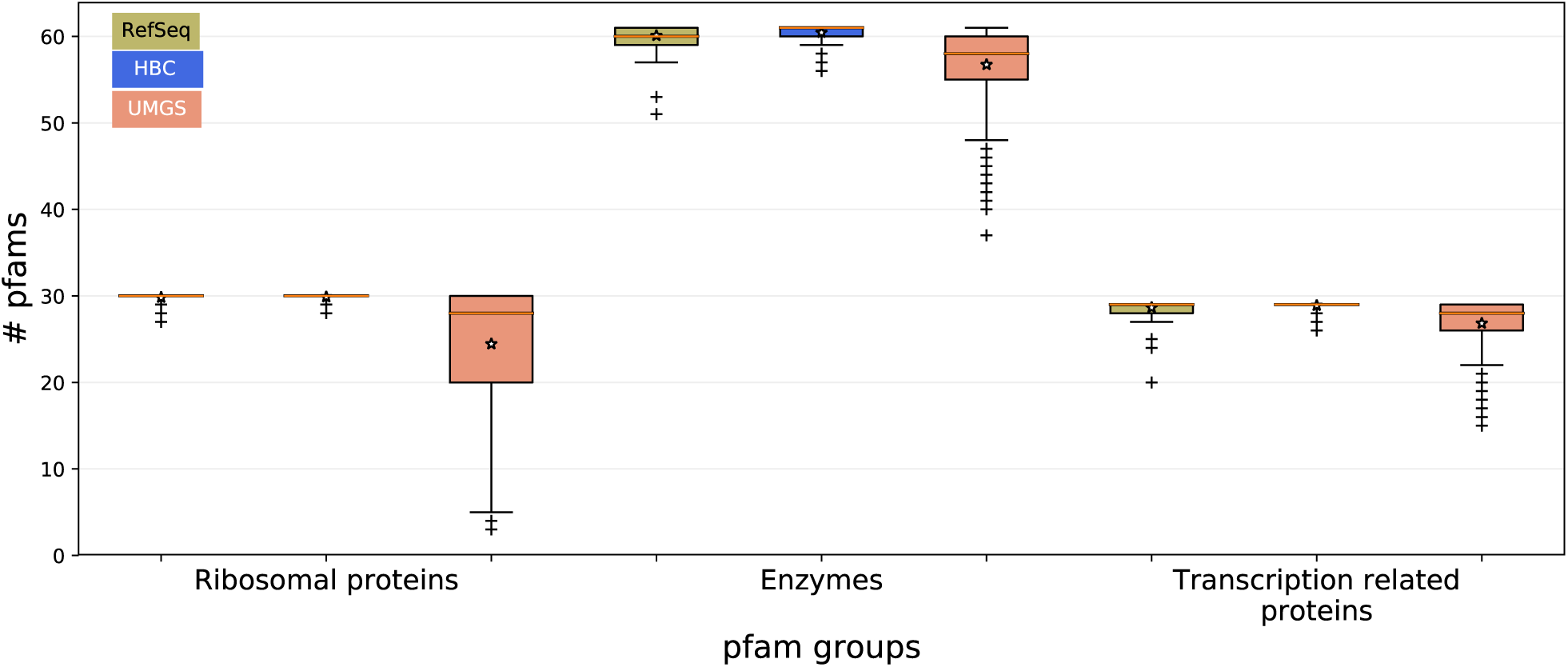
Boxplots showing the distribution of the presence of ribosomal protein, enzyme and transcription related protein pfam markers in genomes three different sources.

**Figure 3:**
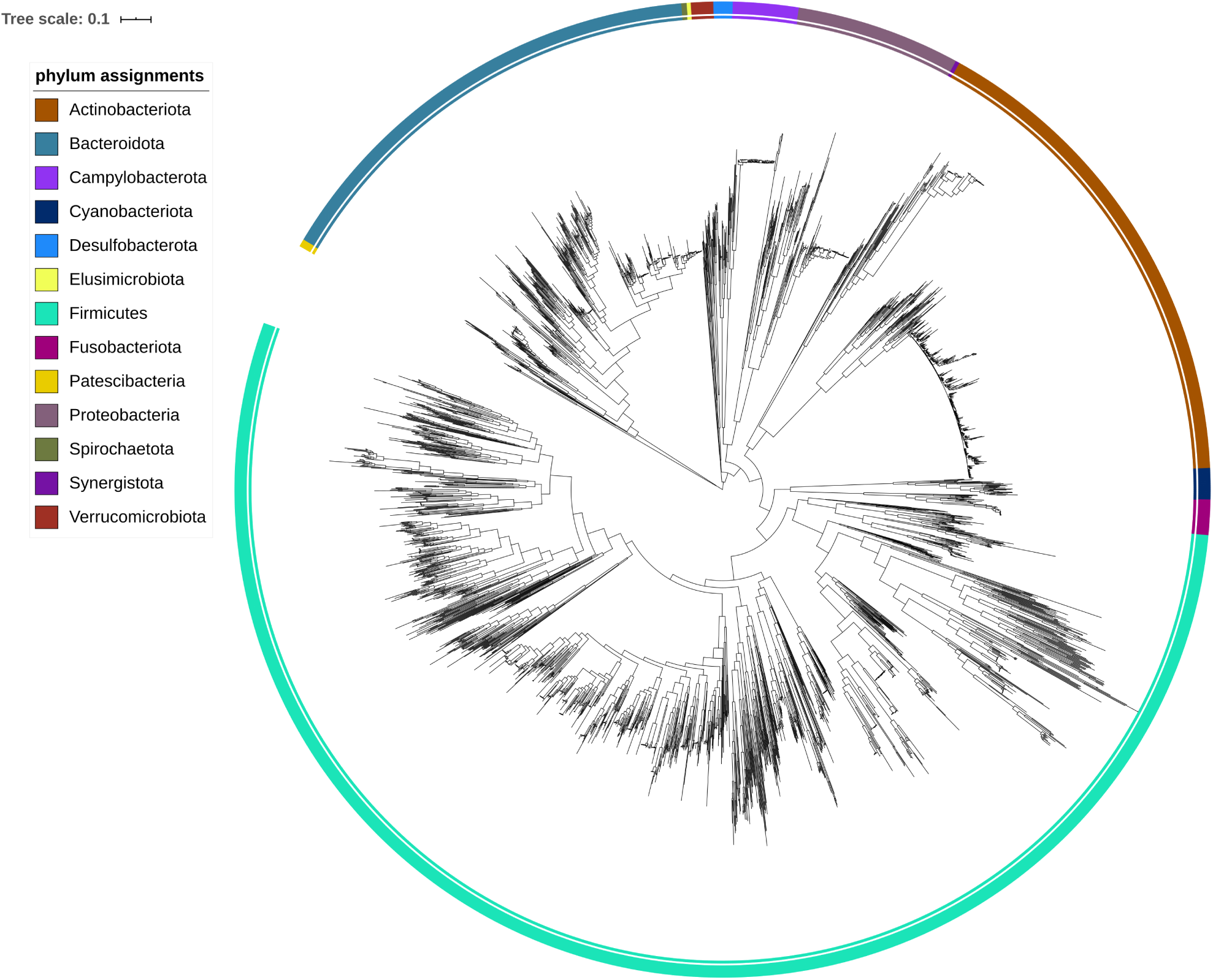
Phylogeny of 3,293 gut genomes, including 2,678 MAGs (736 HBC and 1,943 UGMS), and 614 RefSeq genomes. The phylogenetic tree was constructed from an alignment combining alignments of 62 marker Pfams (see Methods for details). All these genomes were assigned to 16 phyla.

The tree was then used to derive taxonomic annotations for gut species using the least common ancestors approach implemented in GTDB-toolkit^54^. All 3,293 genomes were assigned up to class level taxonomic assignments. These genomes were classified into 13 groups at the phylum level. These phyla are, in the order of their size: *Firmicutes* (1,831 genomes), *Actinobacteria* (550), *Bacteroidota* (511), *Proteobacteria* (184), *Campylobacterota* (74), *Fusobacteriota* (39), *Cyanobacteriota* (35), *Verrucomicrobiota* (25), *Desulfobacterota* (21), *Patescibacteria* (8), *Spirochaetota* (6), *Elusimicrobiota* (5) and *Synergistota* (4). Quantities of the top four phyla is in agreement with previous work^55^. Figure 3 shows the phylum level classifications. A more detailed classification into different taxonomic levels is summarized in Table 1. In total, we were able to classify 1,638 genomes/MAGs at the species level, resulting in a total of 856 unique species. Of the remaining 1,661 bins that lack species level taxonomic assignments, 1,522 were from the UMGS MAGs. Nearly 87% (1,447 out of 1,661) of the taxonomically unassigned MAGs at the species level come from 5 classes, Clostridia, Coriobacteriia, Bacteroidia, Bacilli and Negativicutes, in decreasing order of their contribution. In total, there were 22 class level taxonomic classifications representing all of 3,293 genomes. The distribution of these genomes among these 22 classes is shown in Figure 4. A significant proportion of the genomes belonging to the 5 classes have no species level assignments, especially for the class Coriobacteriia, where most of its genomes could not be identified at the species level. This reflects the diversity of gut bacteria and how some of the taxonomic classes in the current reference databases are underrepresented.

**Figure 4:**
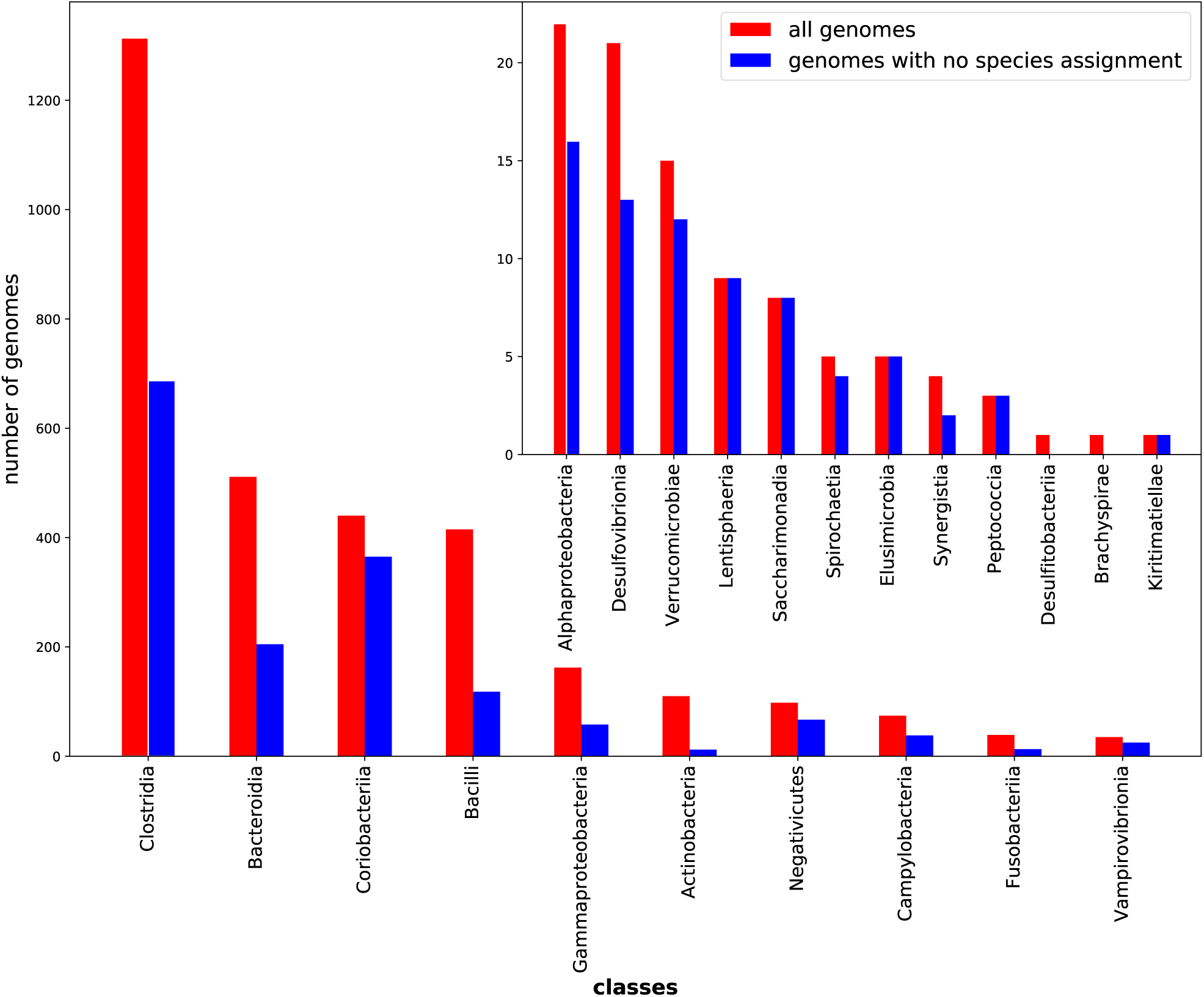
Barplot representing the number of genomes found in the different class level taxonomic classifications. Total number of genomes in each class is given in red, and total number of genomes that do not have species level taxonomic assignments are given in blue. The subplot within the main plot is zoomed in over classes with fewer genomes in them.

**Table 1:** Comparison of the taxonomic distribution of the gut genomes from different resources

| collection | kingdom | phylum | class | order | family | genus | species |
| --- | --- | --- | --- | --- | --- | --- | --- |
| NCBI | 614 | 614 | 614 | 614 | 614 | 614 | 564 |
| HBC | 736 | 736 | 736 | 736 | 734 | 726 | 647 |
| UMGS | 1943 | 1943 | 1943 | 1936 | 1876 | 1554 | 421 |

Our gut tree was in complete agreement with the taxonomies assigned by the GTDB toolkit^54^ up to class level annotations. There were however a few cases of disagreements at the order level between the placement of certain genomes within the tree and the annotations received by them. The major conflict was within the order *Bacilliales*. Our constructed gut tree suggests certain orders to be polyphyletic, in particular the *Lactobacillales* and *Bacillales* clades. Some initiative has already been taken by the authors of GTDB-tk^54^, in dividing the order *Bacillales* into two clades, *Bacillales* and *Bacillales A*. Our tree suggests further divisions of the *Bacillales* clade. Moreover there were two species from the *Lactobacillales* order that do not share the same immediate parent as the remaining members of this order, suggesting the polyphyletic nature of this order as well. These inconsistencies are depicted with detail in Supplementary Figure 3. To understand further the reasons behind these inconsistencies we isolated the problem by focusing on the clade where they arise. The latter clade was the class bacilli, where we had a total of 414 genomes. After extracting this subset of genomes we inferred a phylogeny in a similar fashion as described above and quantified branch support values by a 100 bootstrapped aggregates. The bootstrapped tree is summarized in Supplementary Figure 4. A more focused view over the branches where the splits lead to these inconsistencies alongside their bootstrap supports can be seen in Supplementary Figures 5 and 6. These figures show that the branches leading to these inconsistencies have relatively high support over the bootstrap datasets, with the lowest branch having a support of 89%. Such high support values suggest the polyphyletic nature of *Lactobacillales* and *Bacillales* clades and that the annotation received by the underlying genomes might be prone to some errors. Despite the high support values for these problematic branches, the branch lengths between those splits and their immediate parents are relatively small, indicating certain limitations of the inferred tree and the molecular information arising from the marker genes used to group those species together.

Finally we quantified the distributions of evolutionary distances between genomes at different taxonomic levels. Species pairs were defined for each taxonomic level as pairs of genomes whose most recent common ancestor is at the specified level (i.e. if we’re enumerating class level pairwise relationships, then the common ancestor of any genome pair within this group is at the class level). We used only the genomes/MAGs with species annotation for this analysis. Figure 5 summaries the distribution of phylogenetic distances at different taxonomic relationships, which shows discrete boundaries between the different levels of taxonomic relationships.

**Figure 5:**
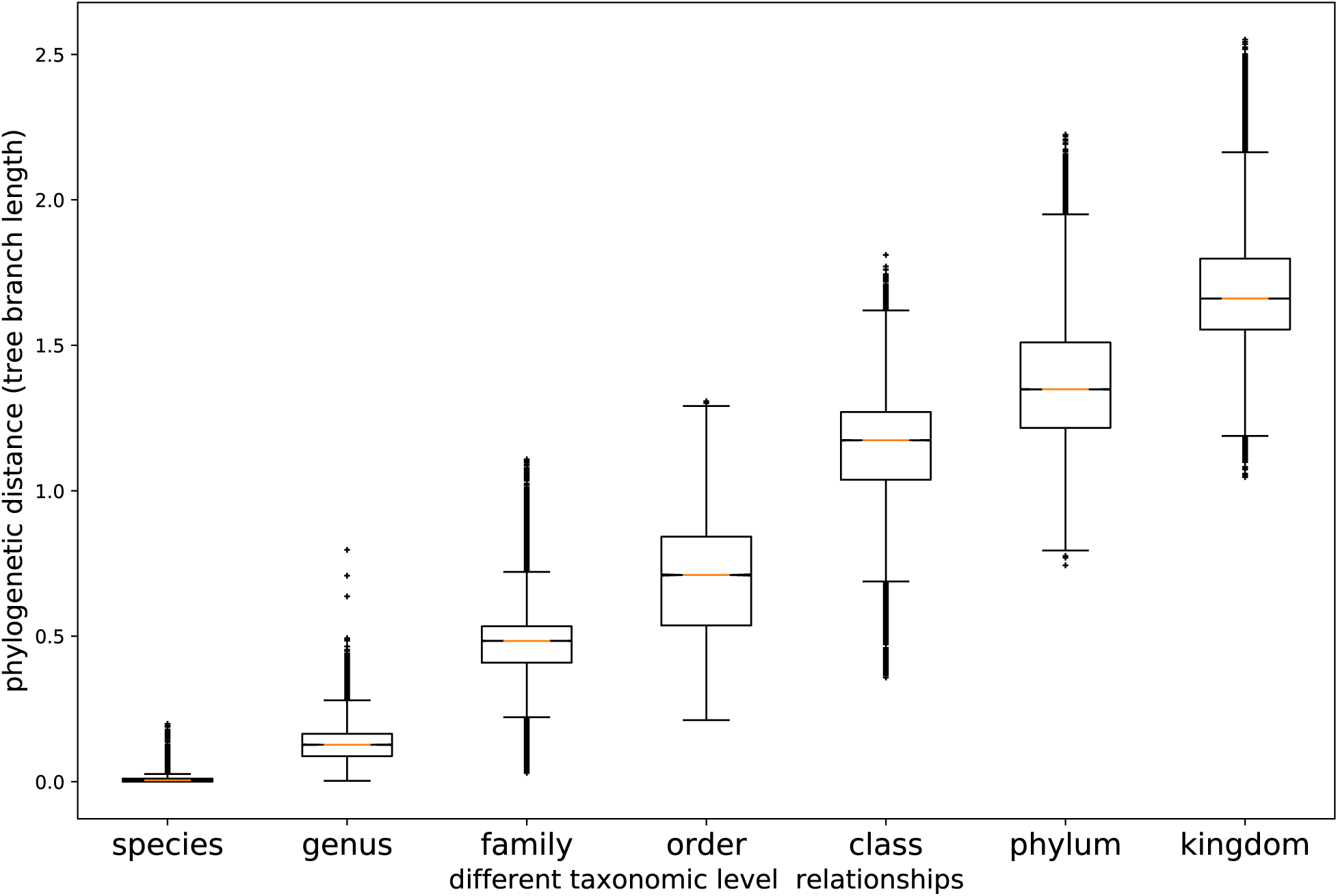
Boxplot representing the distribution of phylogenetic distances between pairs of genomes at different level of evolutionary relations.

### 3.3 Application of the gut tree to peptide-centric metaproteomics studies

Having an annotated comprehensive bacterial species tree, one useful application is to identify prominent species in different metaproteomics samples. For demonstration purposes we used a sample metaproteomics data set from Unipept’s website ^35^, obtained from a human gut through shotgun mass spectrometry ^56^. Since we are only interested in Bacterial species in this study, Eukaryoteic and Archaeal mapped peptides were removed. A total of 1,150 peptides were explained by the species present in the Unipept pipeline; the number of identified peptides increased by 35% when searching the same sample against our bacterial tree, resulting in a total of 1,558 peptides mapped to at least one species within the tree. All of the peptides identified by Unipept were also identified by our tree, in addition to 408 new peptides not identified by the latter. Figure 6 summarises these results, in which we collapsed the clades in our gut tree to “class” level for clarity (links to iTOL^57^ are provided in the Supplementary Material for users to explore the gut tree interactively).

**Figure 6:**
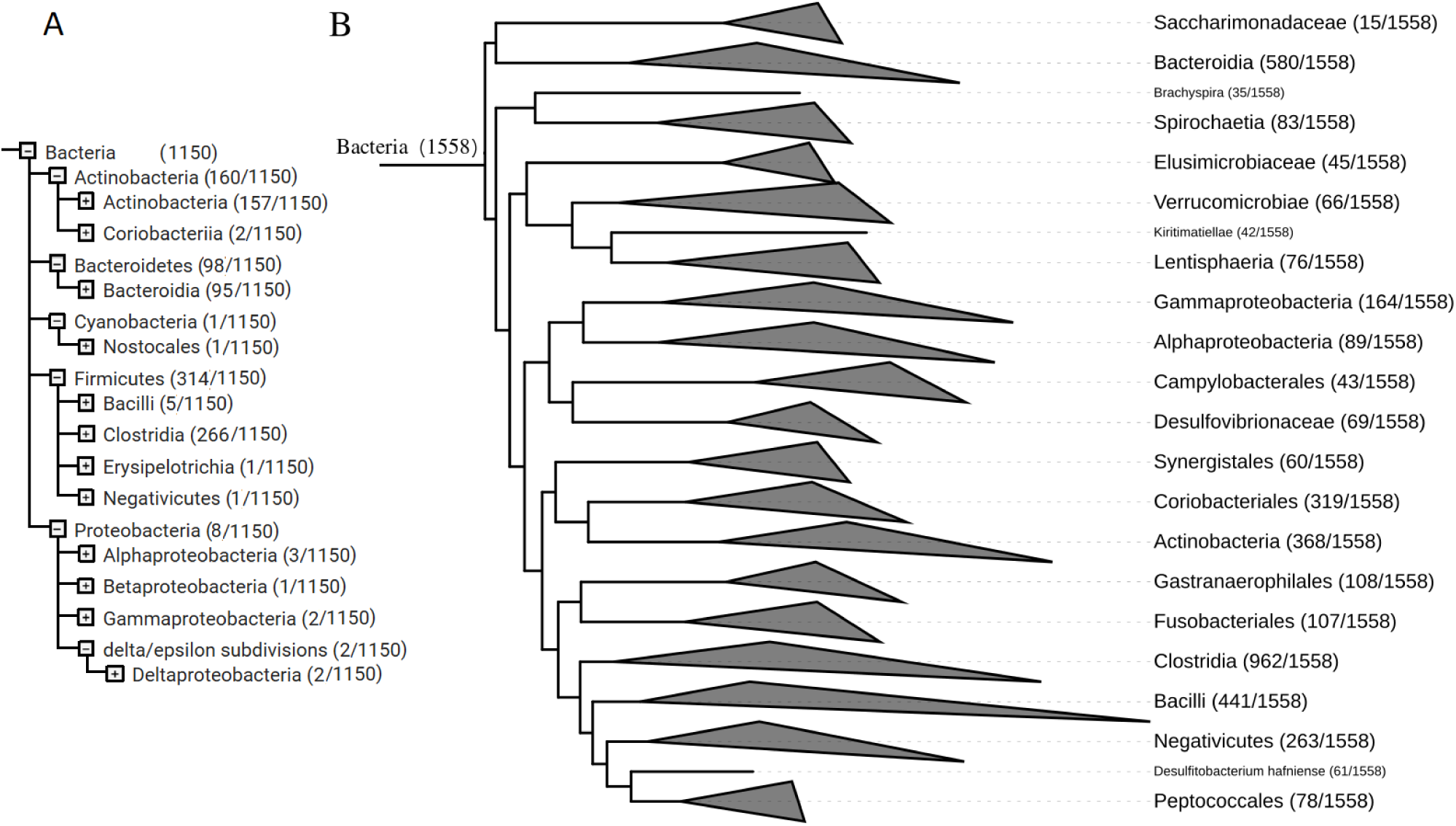
Comparison of the tree coverage of peptides derived from metaproteomics data. Results obtained from Unipept are on the left (A), and those obtained using out tree are to the right (B). Numbers in parentheses indicate the number of peptides mapped to each clade. Clades are collapsed to class level for clarity.

### 3.4 Application of the gut tree to metagenomics studies

Here we demonstrate that the gut tree can be used for taxonomic profiling given a metagenomics dataset. We used this metagenomic dataset^51^ (accession: SRR769523) as an example. We applied Bowtie2 to map the reads onto the big collection of gut bacterial genomes as well as a smaller collection with only 47 representative species ^51^ for comparison. We also applied MetaPhlAn2^52^ and Kraken2^48^ to analyze this dataset. Only 13.80% of the raw reads were mapped to the 47 representative species; by contrast, this number increased (by more than 6 fold) to 85.39% when all genomes on our gut tree were included for reads mapping. Kraken2 mapped 64.1% of the total number of reads (MetaPhlAn2 only considers marker genes for reads mapping so this number is not reported for MetaPhlAn2). Table 2 summarizes the number of unique species identified by each method when different minimum relative abundances were applied. While kraken2 reports an extremely large number of unique species without relative abundance constraint (5,265 species, which is likely an overestimation of the species diversity), all three methods report similar results when only species of minimum relative abundance (0.1% or 1.0%) were included, as seen in Table 2.

**Table 2:** A summary of the number of unique species identified by Metaphlan2, Kraken2 and our gut-tree based approach as a function of different relative abundance cutoffs (metagenomic sample SRR769523).

| Minimum relative abundance (%) | MetaPhlAn2 | kraken2 | Gut-tree (LCA) |
| --- | --- | --- | --- |
| 0 (no filtering) | 73 | 5,265 | 336 |
| 0.1 | 43 | 64 | 78 |
| 1.0 | 21 | 24 | 29 |

We compared the taxonomic profiling results from the different approaches. As MetaPhlAn2 and Kraken2 do not provide a tree visualization, we summarized all of the taxonomic profiling results at the class level for demonstration purpose. Table 3 shows that our approaches and MetaPhlAn2 gave more similar taxonomic profiles as compared to Kraken2. MetaPhlAn2 and our approaches reported similar relative abundances for the top two most abundant classes; the discrepancy of the relative abundances of Bacilli and Erysipelotrichia is a result of the discrepancy of the taxnomic assignments of the genomes (MetaPhlAn2 uses NCBI taxonomy which has Erysipelotrichia as a class, whereas our approaches use GTDB taxonomy which has Erysipelotrichia as an order under the Bacilli class); and for Negativicutes, we note that although the same number of reads were mapped to genomes belonging to Nagativicutes (mostly to *Dialister invisus* and *Veillonella parvula*) by our approach and MetaPhlAn2, our gut tree has expanded collections of genomes in other clades so the relative abundance of this clade is decreased in our analysis. The relative abundance for Bacteroidia reported by Kraken2 was significantly higher than the ones reported by MetaPhlAn2 and our approaches. For Kraken2, we only show the top 20 classes it reported, which explain 98.9% of the mapped reads; we observed that beyond these classes, Kraken2 started to report clades that are likely to be false positives, including Flavobacteria (which is known to be found in marine and freshwater environments) and Mollicutes (previously found in respiratory and urogenital tracts). We note that our gut tree doesn’t include archaea genomes so our analysis doesn’t report the abundance of Methanobacteria.

**Table 3:** A summary of the percentage of reads mapped to different taxonomic units using MetaPhlAn2, Kraken2 and our gut tree based approach (metagenomic sample SRR769523).

| Class | MetaPhlAn2 | kraken2 | Gut-tree (multi-mapped) | Gut-tree (LCA) |
| --- | --- | --- | --- | --- |
| Clostridia | 56.87% | 42.05% | 58.31% | 50.07% |
| Bacteroidia | 24.35% | 48.98% | 25.86% | 38.08% |
| Bacilli | 0.1% | 1.37% | 11.18% | 6.69% |
| Erysipelotrichia | 10.44% | 1.15% | N/A | N/A% |
| Negativicutes | 3.64% | 0.15% | 1.6% | 1.04% |
| Coriobacteriia | N/A | 1.23% | 0.89% | 1.97% |
| Methanobacteria | 2.76% | 1.33% | N/A | N/A% |
| Actinobacteria | 0.92% | 0.81% | 0.26% | 0.10% |
| Deltaproteobacteria | 0.56% | 0.29% | N/A | N/A |
| Betaproteobacteria | 0.23% | 0.29% | N/A | N/A |
| Verrucomicrobiae | 0.07% | 0.30% | 0.75% | 0.78% |
| Alphaproteobacteria | N/A | 0.42% | 0.04% | 0.43% |
| Gammaproteobacteria | 0.02% | 0.69 % | 0.56% | 0.34% |
| Desulfovibrionia | N/A | NA | 0.36% | 0.44% |
| Vampirovibrionia | N/A | NA | 0.04% | 0.03% |
| Spirochaetia | N/A | N/A | 0.08% | 0.005% |
| Saccharimonadia | N/A | N/A | 0.001% | 0.001% |
| Fusobacteriales | N/A | N/A | 0.003% | 0.002% |
| Synergistia | N/A | 0.04 % | 0.001% | 0.003% |
| Lentisphaeria | N/A | N/A | 0.003% | 0.003% |
| Peptococcia | N/A | N/A | 0.001% | 0.001% |
| Flavobacteria | N/A | 0.14% | N/A | N/A |
| Cytophagia | N/A | 0.09% | N/A | N/A |
| Mollicutes | N/A | 0.06% | N/A | N/A |
| Mammalia | N/A | 0.06% | N/A | N/A |
| Epsilonproteobacteria | N/A | 0.05% | N/A | N/A |

Our approaches provide quantification of the species at the different taxonomic levels summarized in a tree, which can be visualized using iTOL^57^ for example. Figure 7 shows the results using the LCA approach to assign reads to the nodes in the gut tree. Supplementary Figure 9 shows the results using the multi-mapped approach for quantification.

**Figure 7:**
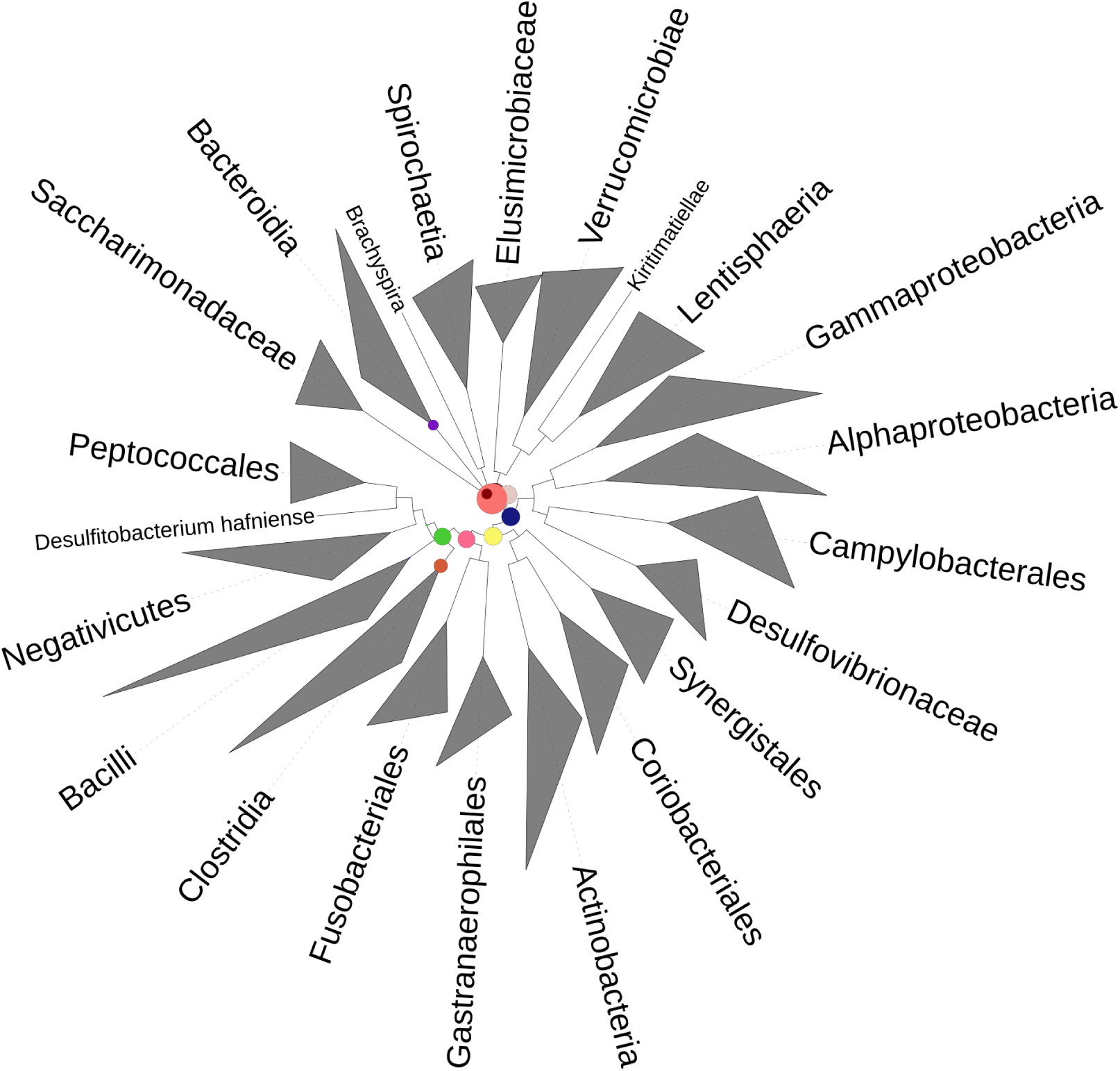
A summary of the species diversity of a metagenome dataset on the gut tree. The circles on the tree represent number of fragments (paired end reads) mapped to each clade using the LCA approach. Their sizes are proportional to the count. Clades are collapsed to class level in this case for clarity.

## 4 Discussion

In this study, we present a comprehensive human gut bacterial tree, aiming to provide a more complete view for metagenomic applications related to human gut, as the ones highlighted here. The majority of the genomes used here were derived computationally as metagenome-based assembled genomes, leading to a certain extent of incompleteness and contamination. We showed that although alignment free methods—developed to cope with such case—had a significantly inferior performance compared to traditional multiple sequence alignment-concatenation-based approaches (Figure 1), when we attempt to cluster these genomes. Our choice of FastTree as an alignment-oriented phylogeny inference tool was purely based on computational considerations. Our final alignment had 3,293 rows and 33,182 columns, which is too large for the more rigorous maximum likelihood and Bayesian-based tree inference methods, including RAxML^30,31^. When comparing FastTree to RAxML over our simulated data set of 400 species, we show both tools gave similar results (Figure 1 (A)). Furthermore when comparing these two tools over a data set with missing genes—which is the case presented here—the results from the two methods correlate almost perfectly (Figure 1 (B)). Similar results were also observed when FastTree was compared to RAxML over large datasets in^54^, suggesting differences resulting from using FastTree as opposed to the computationally more demanding tools can be considered insignificant.

Our annotation of the final gut tree is based on the GTDB-tk annotation, since it provides a consistent annotation, standardized over all genomes. We notice some biases in annotations when comparing proportions of number of genomes found across different taxonomic clades to the number of genomes receiving a complete annotation (up to species level annotation, Figure 4). Certain species classes such as *Clostridia* and *Coriobacteriia* are both relatively large clades and almost entirely annotated up to species levels, whereas there are those such as *Lentisphaeria* and *Saccharimonadia* that are smaller clades with no species level annotations. The latter suggests the presence of a noticeable bias in the scientific community, which has so far only focused on specific parts of the bacterial tree of life, leaving a large proportion yet to be uncovered. With the increase in sequencing throughput and culture free binning methods becoming more and more standardized we should expect to shed more light over the microbial dark matter in the near future.

Although our gut tree is significantly smaller than that presented by the GTDB-toolkit, it contains more gut bacteria. We note that the majority of the genomes participating in the gut tree are novel, and therefore many of the novel bacteria didn’t receive family to species level annotation by GTDB-tk, but they are still useful for presenting the diversity and complexity of human gut microbiome. We observed some disagreements between the inferred taxonomies using GTDB-tk and the inferred phylogeny using our tree, and we think it is important to report these disagreements so users can be aware of them.

We note that a tree with expanded list of bacteria species, many unseen, may suggest evolutionary relationship that disagree with previously proposed taxonomic assignments for some species. Polyphyletic clades were proposed to deal with such cases, for example, the order *Bacillales* was divided into two clades, *Bacillales* and *Bacillales A*^54^. Our tree suggests further divisions of the *Bacillales* clade. Moreover there were two species from the *Lactobacillales* order that do not share the same immediate parent as the remaining members of this order, suggesting the polyphyletic nature of this order as well. We believe that changes to taxonomic annotations more drastic than proposing polyphyletic clades will be needed in the future when even more diversity of species are expected. Nevertheless, we showed that using a diverse and comprehensive tree helps improve the analysis of metaproteomics and metagenomics data, as demonstrated in the two applications we presented.

## 5 Conclusions

In this study we constructed a comprehensive human gut bacterial tree, containing a large number of metagenome assembled genomes (MAGs), and developed two downstream applications of the tree for metagenomic and metaproteomic data analysis. We showed that the performance of RAxML, which is usually regarded as a more profound and accurate tree inferring tool compared to FastTree, was comparable to that of the latter when we inferred trees using a relatively large set of species. Including MAGs in the tree provides more comprehensive representation of species diversity associated with human gut microbiome, important for studying the taxonomic composition of gut microbiome. Furthermore we provide an automated pipeline allowing users to infer phylogeny starting from any set of genomes, MAGs or both.

## Declarations

### Ethics approval and consent to participate

Not applicable

### Consent for publication

Not applicable

### Availability of data and material

The gut tree and companion tools are available at https://github.com/mgtools/guttree and http://omics.informatics.indiana.edu/guttree/.

## Competing interests

The authors declare that they have no competing interests.

### Funding

The NIH grants 1R01AI108888 and 1R01AI143254.

### Authors’ contribution

All authors contributed equally in designing and performing the experiments, writing and revising the manuscript.

## Acknowledgements

This research was partially supported by the Indiana University (IU) Precision Health Initiative (PHI).

## References

[1] Hug, L. A., Baker, B. J., Anantharaman, K., Brown, C. T., Probst, A. J., Castelle, C. J., Butterfield, C. N., Hernsdorf, A. W., Amano, Y., Ise, K., Suzuki, Y., Dudek, N., Relman, D. A., Finstad, K. M., Amundson, R., Thomas, B. C., and Banfield, J. F. (Apr, 2016) A new view of the tree of life. Nat Microbiol, 1, 16048.

[2] Woese, C. R., Kandler, O., and Wheelis, M. L. (1990) Towards a natural system of organisms: proposal for the domains Archaea, Bacteria, and Eucarya.. Proceedings of the National Academy of Sciences, 87(12), 4576–4579.

[3] Fleischmann, R. D., Adams, M. D., White, O., Clayton, R. A., Kirkness, E. F., Kerlavage, A. R., Bult, C. J., Tomb, J.-F., Dougherty, B. A., Merrick, J. M., et al. (1995) Whole-genome random sequencing and assembly of Haemophilus influenzae Rd. Science, 269(5223), 496–512.

[4] Schuster, S. C. (2007) Next-generation sequencing transforms today’s biology. Nature methods, 5(1), 16.

[5] Dick, G. J., Andersson, A. F., Baker, B. J., Simmons, S. L., Thomas, B. C., Yelton, A. P., and Banfield, J. F. (2009) Community-wide analysis of microbial genome sequence signatures. Genome biology, 10(8), R85.

[6] Zerbino, D. R. and Birney, E. (2008) Velvet: algorithms for de novo short read assembly using de Bruijn graphs. Genome research, 18(5), 821–829.

[7] Li, D., Liu, C.-M., Luo, R., Sadakane, K., and Lam, T.-W. (2015) MEGAHIT: an ultra-fast single-node solution for large and complex metagenomics assembly via succinct de Bruijn graph. Bioinformatics, 31(10), 1674–1676.

[8] Wu, Y.-W., Tang, Y.-H., Tringe, S. G., Simmons, B. A., and Singer, S. W. (2014) MaxBin: an automated binning method to recover individual genomes from metagenomes using an expectation-maximization algorithm. Microbiome, 2(1), 26.

[9] Imelfort, M., Parks, D., Woodcroft, B. J., Dennis, P., Hugenholtz, P., and Tyson, G. W. (2014) GroopM: an automated tool for the recovery of population genomes from related metagenomes. PeerJ, 2, e603.

[10] Lu, Y. Y., Chen, T., Fuhrman, J. A., and Sun, F. (2017) COCACOLA: binning metagenomic contigs using sequence COmposition, read CoverAge, CO-alignment and paired-end read LinkAge. Bioinformatics, 33(6), 791–798.

[11] Alneberg, J., Bjarnason, B. S., De Bruijn, I., Schirmer, M., Quick, J., Ijaz, U. Z., Lahti, L., Loman, N. J., Andersson, A. F., and Quince, C. (2014) Binning metagenomic contigs by coverage and composition. Nature methods, 11(11), 1144.

[12] Kang, D. D., Froula, J., Egan, R., and Wang, Z. (2015) MetaBAT, an efficient tool for accurately reconstructing single genomes from complex microbial communities. PeerJ, 3, e1165.

[13] Parks, D. H., Imelfort, M., Skennerton, C. T., Hugenholtz, P., and Tyson, G. W. (2015) CheckM: assessing the quality of microbial genomes recovered from isolates, single cells, and metagenomes. Genome research, 25(7), 1043–1055.

[14] Albertsen, M., Hugenholtz, P., Skarshewski, A., Nielsen, K. L., Tyson, G. W., and Nielsen, P. H. (2013) Genome sequences of rare, uncultured bacteria obtained by differential coverage binning of multiple metagenomes. Nature biotechnology, 31(6), 533.

[15] Pasolli, E., Asnicar, F., Manara, S., Zolfo, M., Karcher, N., Armanini, F., Beghini, F., Manghi, P., Tett, A., Ghensi, P., et al. (2019) Extensive unexplored human microbiome diversity revealed by over 150,000 genomes from metagenomes spanning age, geography, and lifestyle. Cell, 176(3), 649–662.

[16] Ferretti, P., Pasolli, E., Tett, A., Asnicar, F., Gorfer, V., Fedi, S., Armanini, F., Truong, D. T., Manara, S., Zolfo, M., et al. (2018) Mother-to-infant microbial transmission from different body sites shapes the developing infant gut microbiome. Cell host & microbe, 24(1), 133–145.

[17] Bäckhed, F., Roswall, J., Peng, Y., Feng, Q., Jia, H., Kovatcheva-Datchary, P., Li, Y., Xia, Y., Xie, H., Zhong, H., et al. (2015) Dynamics and stabilization of the human gut microbiome during the first year of life. Cell host & microbe, 17(5), 690–703.

[18] Bull, M. J. and Plummer, N. T. (2014) Part 1: The human gut microbiome in health and disease. Integrative Medicine: A Clinician’s Journal, 13(6), 17.

[19] Guinane, C. M. and Cotter, P. D. (2013) Role of the gut microbiota in health and chronic gastrointestinal disease: understanding a hidden metabolic organ. Therapeutic advances in gastroenterology, 6(4), 295–308.

[20] Barcenilla, A., Pryde, S. E., Martin, J. C., Duncan, S. H., Stewart, C. S., Henderson, C., and Flint, H. J. (2000) Phylogenetic relationships of butyrate-producing bacteria from the human gut. Appl. Environ. Microbiol., 66(4), 1654–1661.

[21] Pruitt, R. N. and Lacy, D. B. (2012) Toward a structural understanding of Clostridium difficile toxins A and B. Frontiers in cellular and infection microbiology, 2, 28.

[22] Sorg, J. A. and Sonenshein, A. L. (2008) Bile salts and glycine as cogerminants for Clostridium difficile spores. Journal of bacteriology, 190(7), 2505–2512.

[23] Ley, R. E., Turnbaugh, P. J., Klein, S., and Gordon, J. I. (2006) Microbial ecology: human gut microbes associated with obesity. nature, 444(7122), 1022.

[24] Russell, S. L. and Finlay, B. B. (2012) The impact of gut microbes in allergic diseases. Current opinion in gastroenterology, 28(6), 563–569.

[25] Collins, S. M., Surette, M., and Bercik, P. (2012) The interplay between the intestinal microbiota and the brain. Nature Reviews Microbiology, 10(11), 735.

[26] Zielezinski, A., Vinga, S., Almeida, J., and Karlowski, W. M. (10, 2017) Alignment-free sequence comparison: benefits, applications, and tools. Genome Biol., 18(1), 186.

[27] Chan, C. X., Bernard, G., Poirion, O., Hogan, J. M., and Ragan, M. A. (2014) Inferring phylogenies of evolving sequences without multiple sequence alignment. Scientific reports, 4, 6504.

[28] Bernard, G., Ragan, M. A., and Chan, C. X. (2016) Recapitulating phylogenies using k-mers: from trees to networks. F1000Research, 5.

[29] Lu, Y. Y., Tang, K., Ren, J., Fuhrman, J. A., Waterman, M. S., and Sun, F. (2017) CAFE: a C celerated A lignment-F r E e sequence analysis. Nucleic acids research, 45(W1), W554–W559.

[30] Zhou, X., Shen, X.-X., Hittinger, C. T., and Rokas, A. (2018) Evaluating fast maximum likelihood-based phylogenetic programs using empirical phylogenomic data sets. Molecular biology and evolution, 35(2), 486–503.

[31] Liu, K., Linder, C. R., and Warnow, T. (2011) RAxML and FastTree: comparing two methods for large-scale maximum likelihood phylogeny estimation. PloS one, 6(11).

[32] Forster, S. C., Kumar, N., Anonye, B. O., Almeida, A., Viciani, E., Stares, M. D., Dunn, M., Mkandawire, T. T., Zhu, A., Shao, Y., et al. (2019) A human gut bacterial genome and culture collection for improved metagenomic analyses. Nature biotechnology, 37(2), 186.

[33] Almeida, A., Mitchell, A. L., Boland, M., Forster, S. C., Gloor, G. B., Tarkowska, A., Lawley, T. D., and Finn, R. D. (2019) A new genomic blueprint of the human gut microbiota. Nature, p. 1.

[34] Riffle, M., May, D. H., Timmins-Schiffman, E., Mikan, M. P., Jaschob, D., Noble, W. S., and Nunn, B. L. (Dec, 2017) MetaGOmics: A Web-Based Tool for Peptide-Centric Functional and Taxonomic Analysis of Metaproteomics Data. Proteomes, 6(1).

[35] Gurdeep Singh, R., Tanca, A., Palomba, A., Van der Jeugt, F., Verschaffelt, P., Uzzau, S., Martens, L., Dawyndt, P., and Mesuere, B. (Feb, 2019) Unipept 4.0: Functional Analysis of Metaproteome Data. J. Proteome Res., 18(2), 606–615.

[36] Price, M. N., Dehal, P. S., and Arkin, A. P. (2010) FastTree 2–approximately maximum-likelihood trees for large alignments. PloS one, 5(3), e9490.

[37] Parks, D. H., Rinke, C., Chuvochina, M., Chaumeil, P.-A., Woodcroft, B. J., Evans, P. N., Hugenholtz, P., and Tyson, G. W. (2017) Recovery of nearly 8,000 metagenome-assembled genomes substantially expands the tree of life. Nature microbiology, 2(11), 1533–1542.

[38] Jain, R., Rivera, M. C., and Lake, J. A. (1999) Horizontal gene transfer among genomes: the complexity hypothesis. Proceedings of the National Academy of Sciences, 96(7), 3801–3806.

[39] Turnbaugh, P. J., Ley, R. E., Hamady, M., Fraser-Liggett, C. M., Knight, R., and Gordon, J. I. (2007) The human microbiome project. Nature, 449(7164), 804.

[40] Rho, M., Tang, H., and Ye, Y. (2010) FragGeneScan: predicting genes in short and error-prone reads. Nucleic acids research, 38(20), e191–e191.

[41] Finn, R. D., Bateman, A., Clements, J., Coggill, P., Eberhardt, R. Y., Eddy, S. R., Heger, A., Hetherington, K., Holm, L., Mistry, J., et al. (2014) Pfam: the protein families database. Nucleic acids research, 42(D1), D222–D230.

[42] Haft, D. H., Selengut, J. D., and White, O. (2003) The TIGRFAMs database of protein families. Nucleic acids research, 31(1), 371–373.

[43] Finn, R. D., Clements, J., Arndt, W., Miller, B. L., Wheeler, T. J., Schreiber, F., Bateman, A., and Eddy, S. R. (2015) HMMER web server: 2015 update. Nucleic acids research, 43(W1), W30–W38.

[44] Talavera, G. and Castresana, J. (2007) Improvement of phylogenies after removing divergent and ambiguously aligned blocks from protein sequence alignments. Systematic biology, 56(4), 564–577.

[45] Paradis, E., Claude, J., and Strimmer, K. (2004) APE: analyses of phylogenetics and evolution in R language. Bioinformatics, 20(2), 289–290.

[46] Smith, M. R. (2019) Bayesian and parsimony approaches reconstruct informative trees from simulated morphological datasets. Biology letters, 15(2), 20180632.

[47] Felsenstein, J. (1993) PHYLIP (phylogeny inference package), version 3.5 c, Joseph Felsenstein., .

[48] Wood, D. E., Lu, J., and Langmead, B. (2019) Improved metagenomic analysis with Kraken 2. Genome biology, 20(1), 257.

[49] Huson, D. H., Auch, A. F., Qi, J., and Schuster, S. C. (2007) MEGAN analysis of metagenomic data. Genome research, 17(3), 377–386.

[50] Qin, N., Yang, F., Li, A., Prifti, E., Chen, Y., Shao, L., Guo, J., Le Chatelier, E., Yao, J., Wu, L., et al. (2014) Alterations of the human gut microbiome in liver cirrhosis. Nature, 513(7516), 59–64.

[51] Franzosa, E. A., Morgan, X. C., Segata, N., Waldron, L., Reyes, J., Earl, A. M., Giannoukos, G., Boylan, M. R., Ciulla, D., Gevers, D., et al. (2014) Relating the metatranscriptome and metagenome of the human gut. Proceedings of the National Academy of Sciences, 111(22), E2329–E2338.

[52] Truong, D. T., Franzosa, E. A., Tickle, T. L., Scholz, M., Weingart, G., Pasolli, E., Tett, A., Huttenhower, C., and Segata, N. (2015) MetaPhlAn2 for enhanced metagenomic taxonomic profiling. Nature methods, 12(10), 902.

[53] Gurdeep, R. S., Tanca, A., Palomba, A., der Jeugt Van, F., Verschaffelt, P., Uzzau, S., Martens, L., Dawyndt, P., and Mesuere, B. (2019) Unipept 4.0: Functional Analysis of Metaproteome Data.. Journal of proteome research, 18(2), 606–615.

[54] Parks, D. H., Chuvochina, M., Waite, D. W., Rinke, C., Skarshewski, A., Chaumeil, P.-A., and Hugenholtz, P. (2018) A standardized bacterial taxonomy based on genome phylogeny substantially revises the tree of life. Nature biotechnology,.

[55] Khanna, S. and Tosh, P. K. (2014) A clinician’s primer on the role of the microbiome in human health and disease. In Mayo Clinic Proceedings Elsevier Vol. 89, pp. 107–114.

[56] Verberkmoes, N. C., Russell, A. L., Shah, M., Godzik, A., Rosenquist, M., Halfvarson, J., Lefsrud, M. G., Apajalahti, J., Tysk, C., Hettich, R. L., et al. (2009) Shotgun metaproteomics of the human distal gut microbiota. The ISME journal, 3(2), 179.

[57] Letunic, I. and Bork, P. (2019) Interactive Tree Of Life (iTOL) v4: recent updates and new developments. Nucleic acids research, 47(W1), W256–W259.

