## Supplementary file for "A Tree of Human Gut Bacterial Species and its Applications to Metagenomics and Metaproteomics Data Analysis"

### Supporting Information

iTOL tree links

- Link to the Human Gut Bacterial Tree:
  https://itol.embl.de/tree/1565615933232321579123499
- Link to the Human Gut Bacterial Tree Collapsed at the order level:
  https://itol.embl.de/tree/1565615933210131579120606

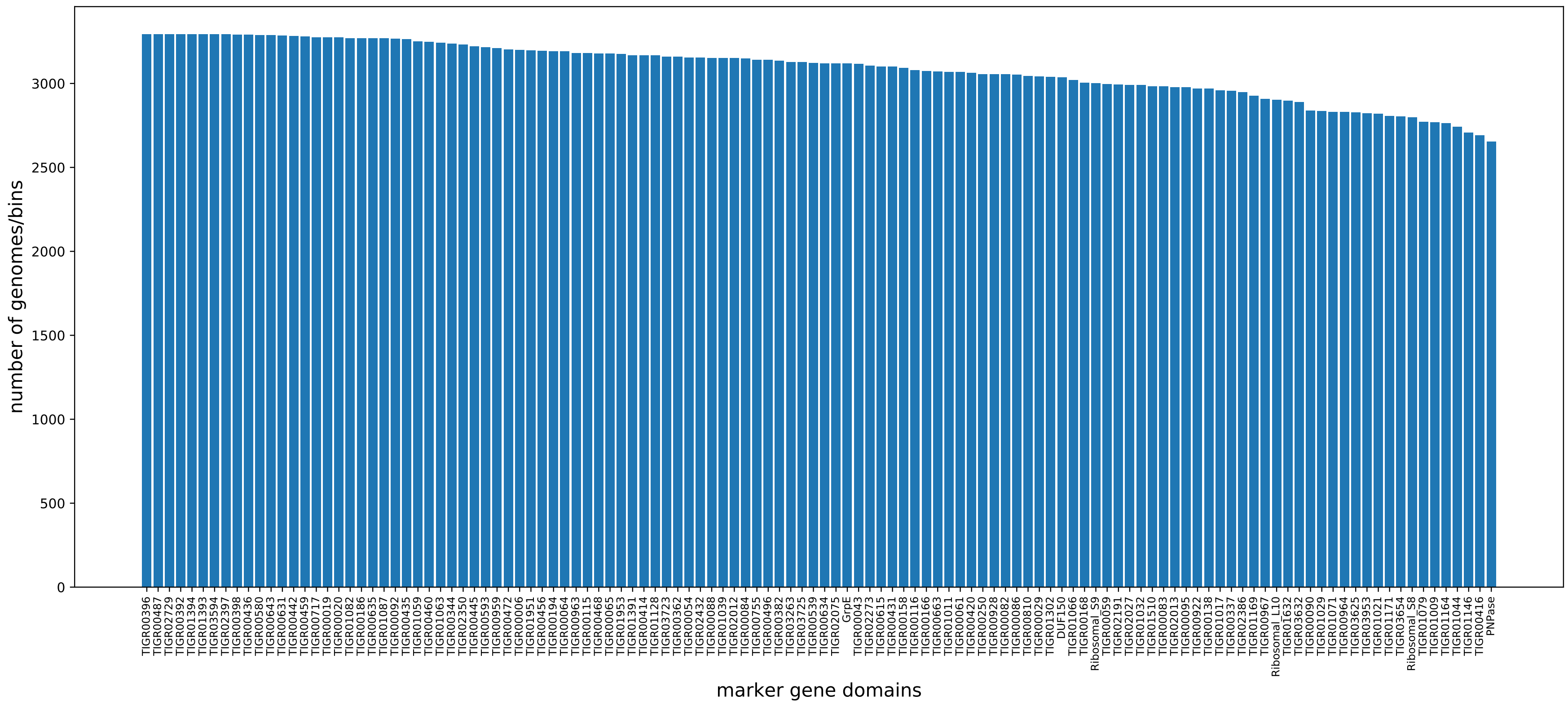


Figure S1: Barplots representing the number of genomes containing the different domains.

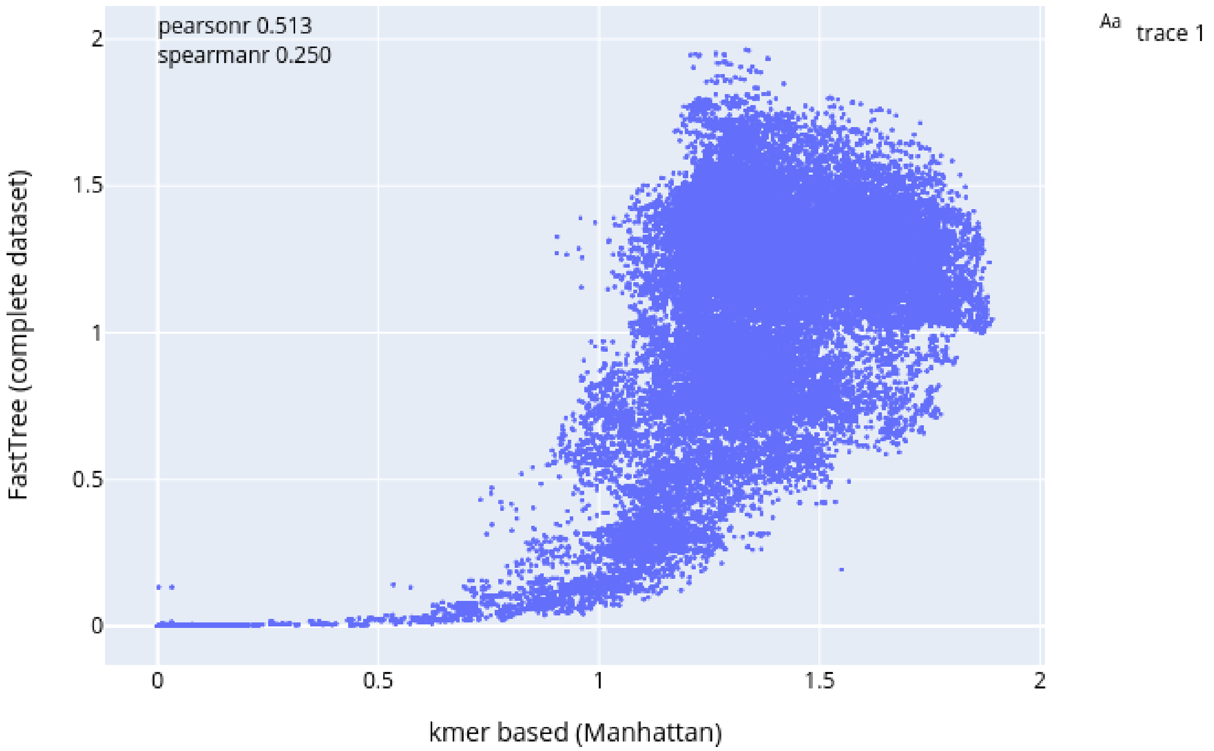


Figure S2: scatter plot showing the correlation of pairwise species distances between alignment free method (using Manhattan distance) and the reference tree constructed for the selected 400 RefSeq genomes.

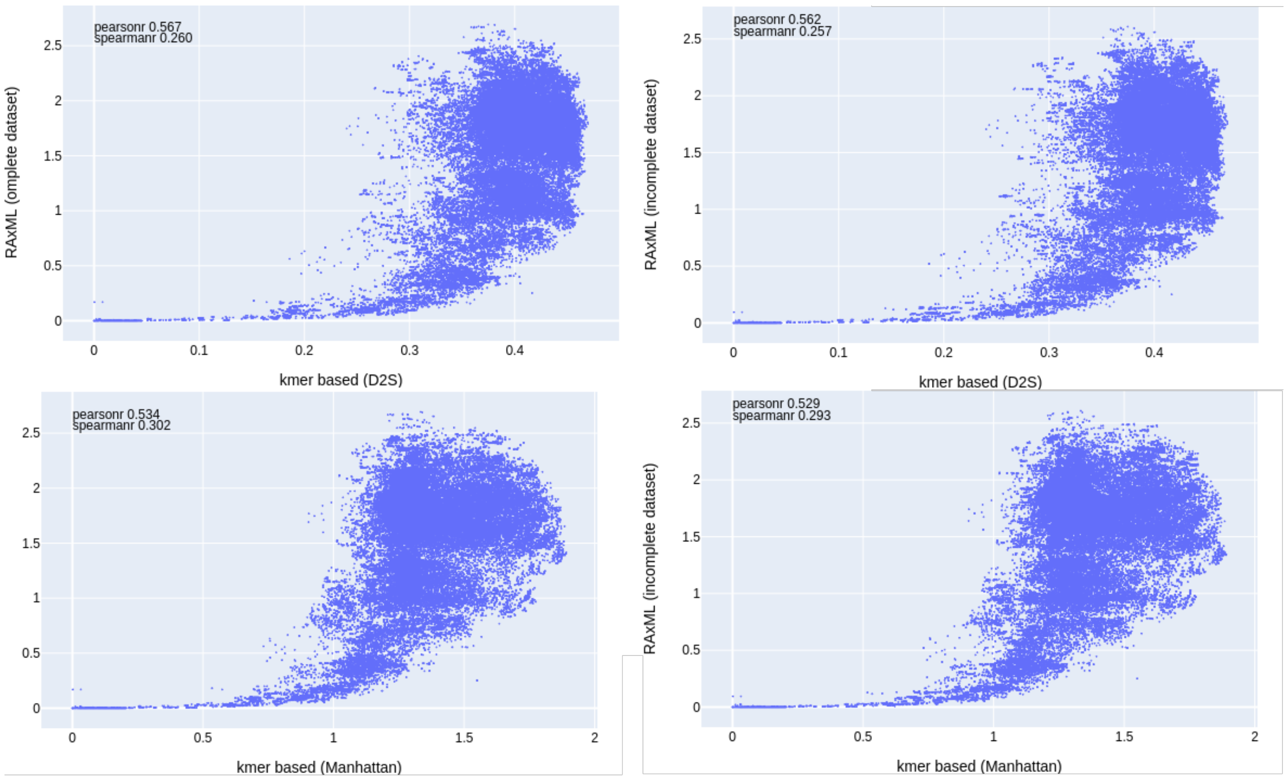


Figure S3: scatter plot showing the correlation of pairwise species distances between alignment free method (using both D2S and Manhattan distances) and the reference tree constructed, using RAxML, for the selected 400 RefSeq genomes.

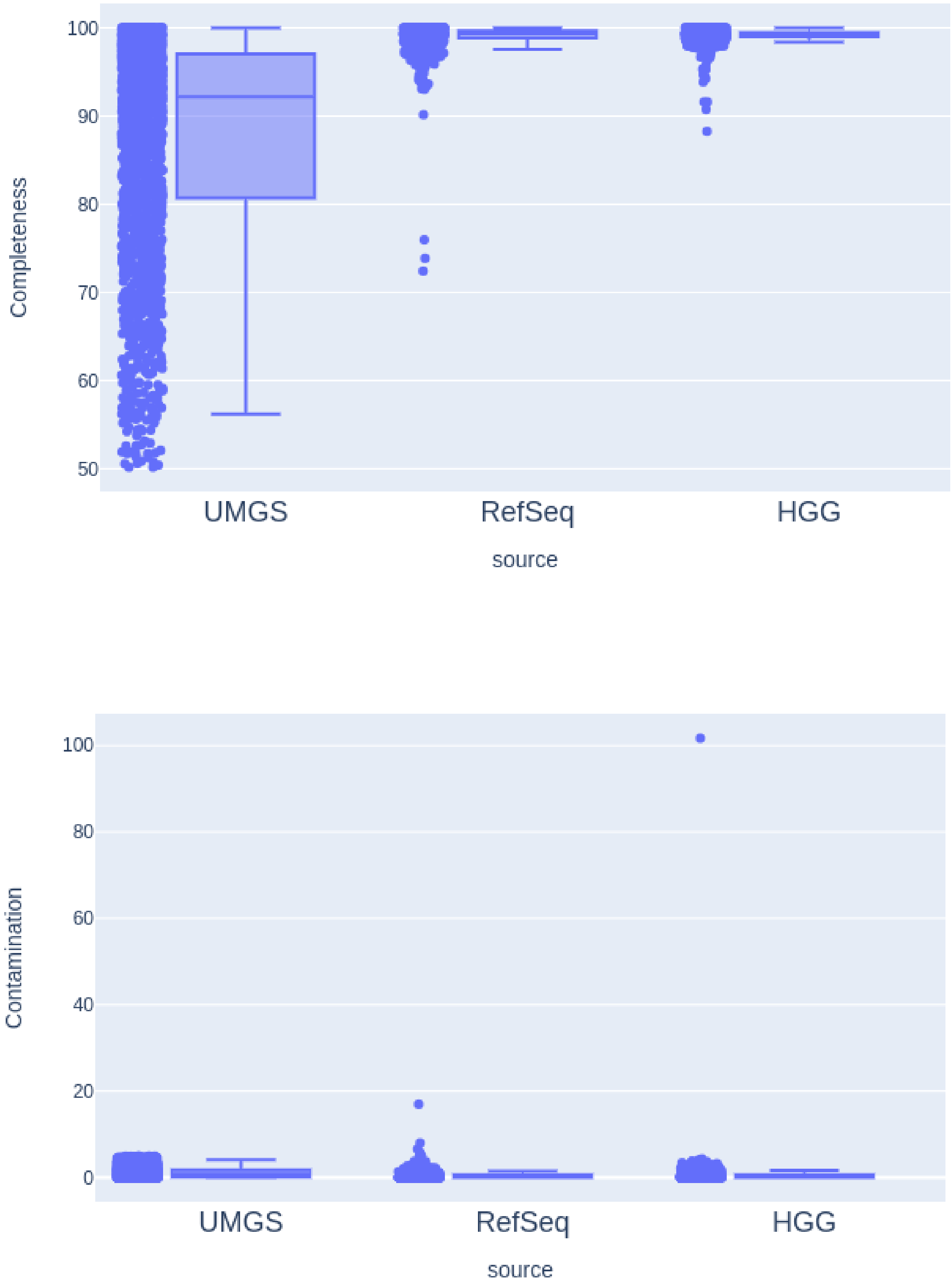


Figure S4: Boxplots summarizing checkm completeness and contamination levels of the bins used to construct the gut-tree

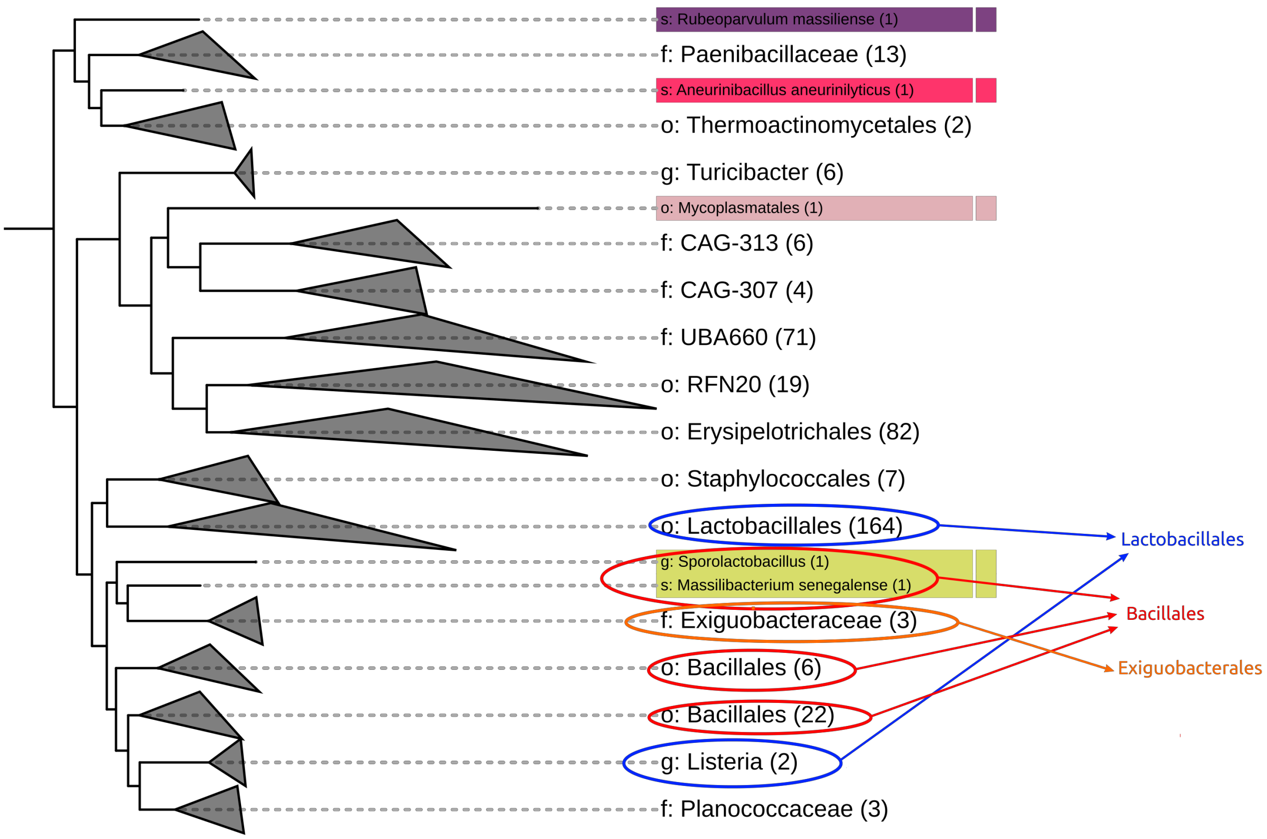


Figure S5: Sub-tree of the human gut tree focusing on the Bacilli class highlighting some inconsistencies.

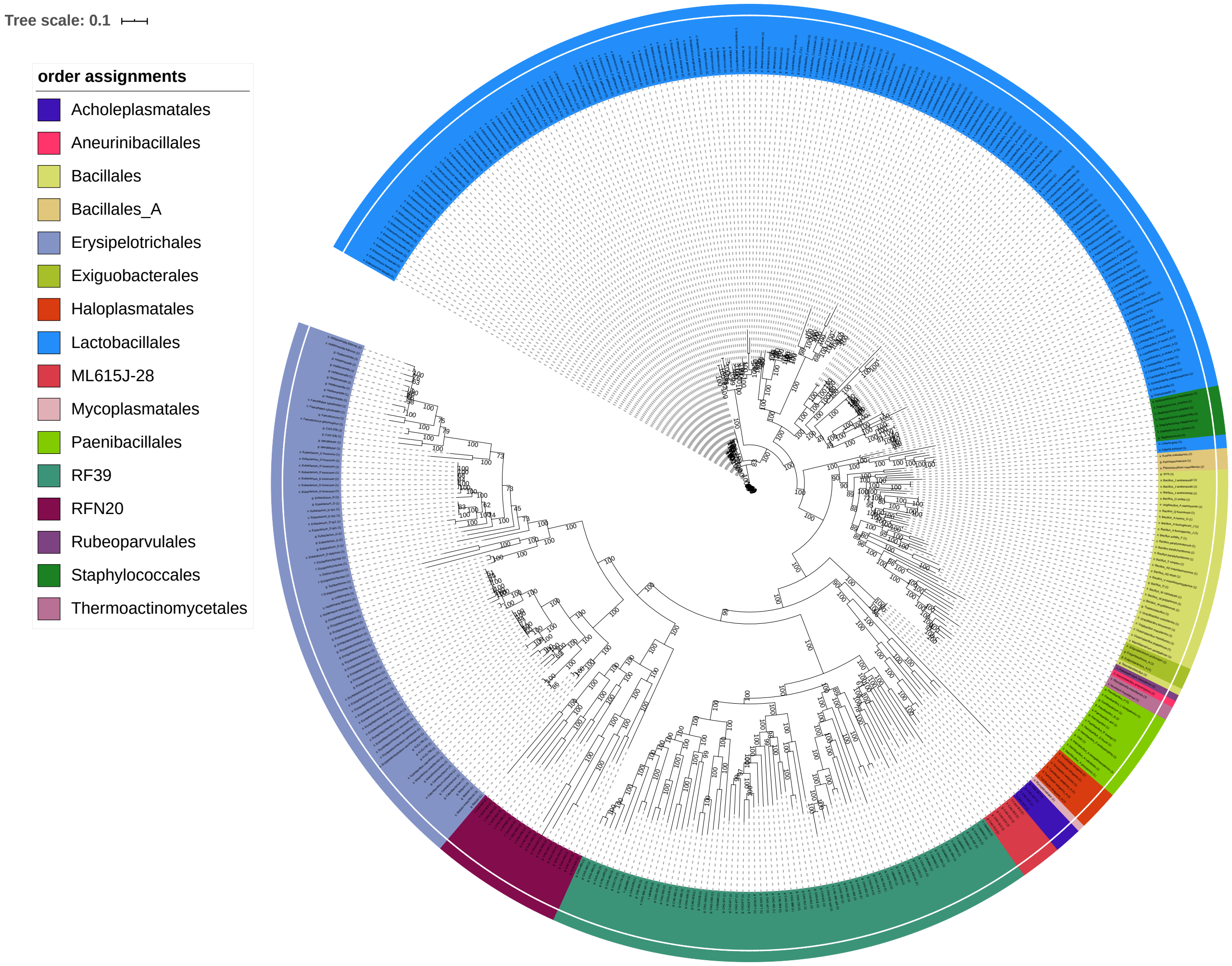


Figure S6: Sub-tree of the human gut tree focusing on the Bacilli with branch support values.

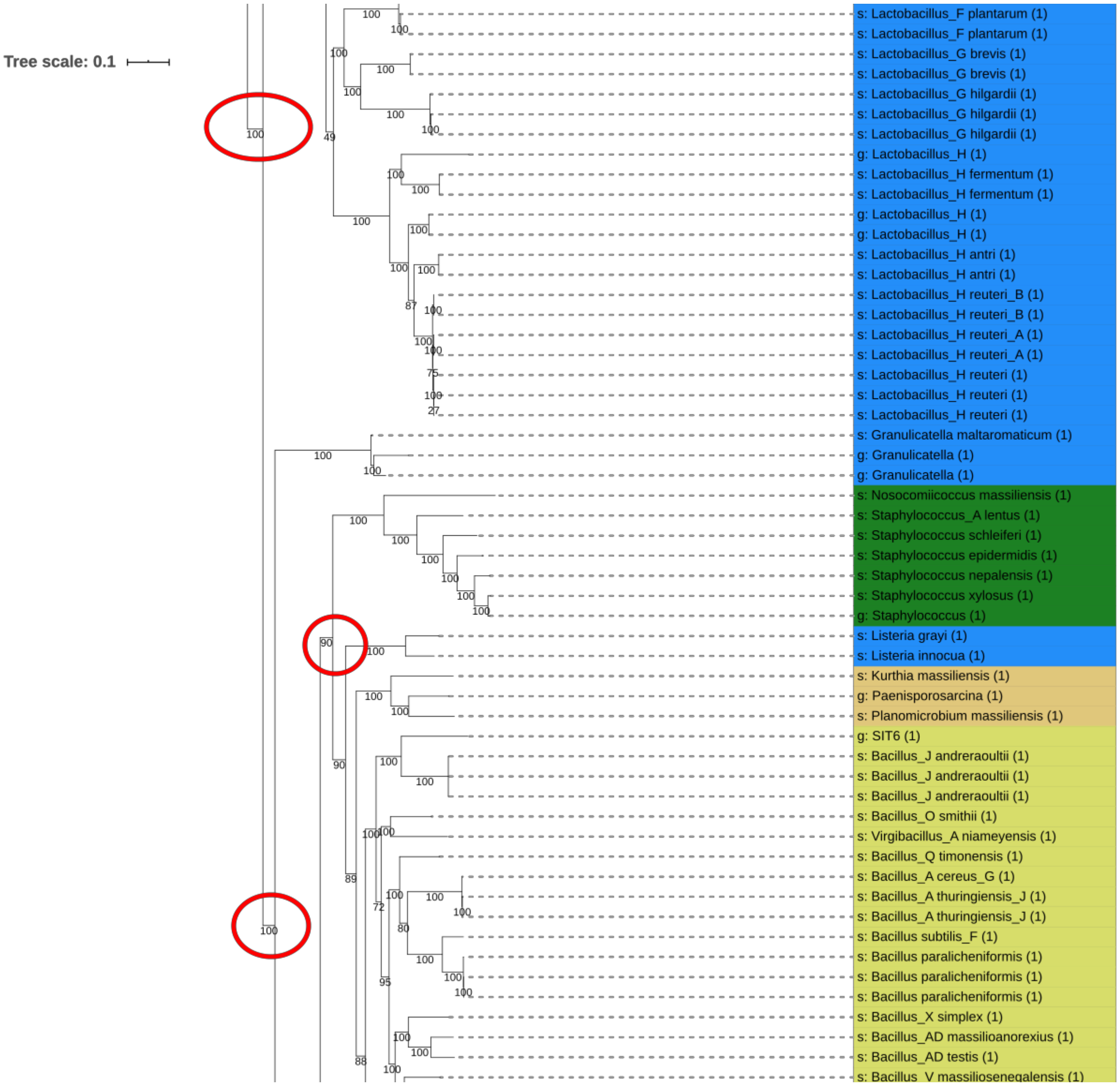


Figure S7: Subsection I of the Bacilli clade, over branches with inconsistent splits with branch support values.

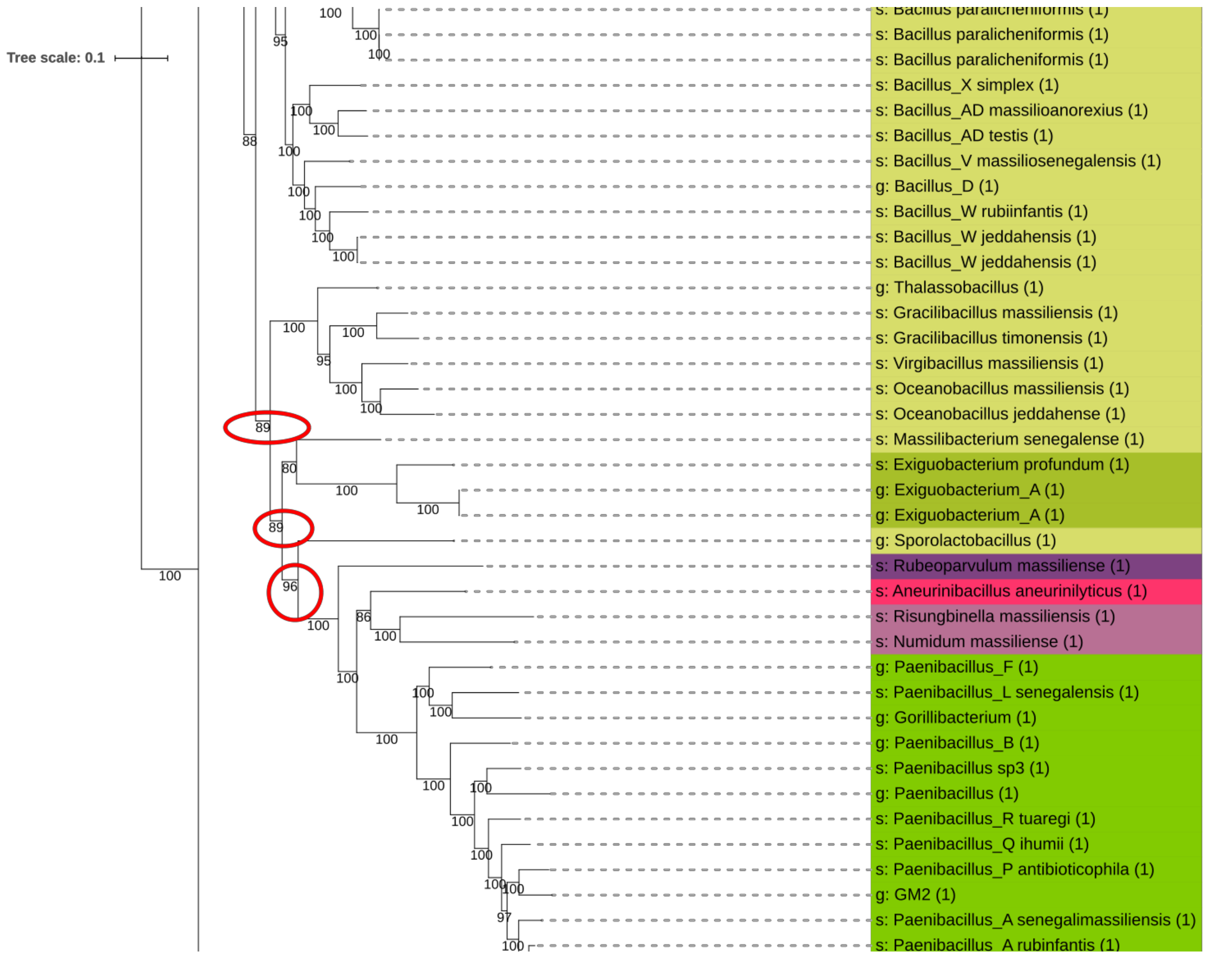


Figure S8: Subsection II of the Bacilli clade, over branches with inconsistent splits with branch support values.

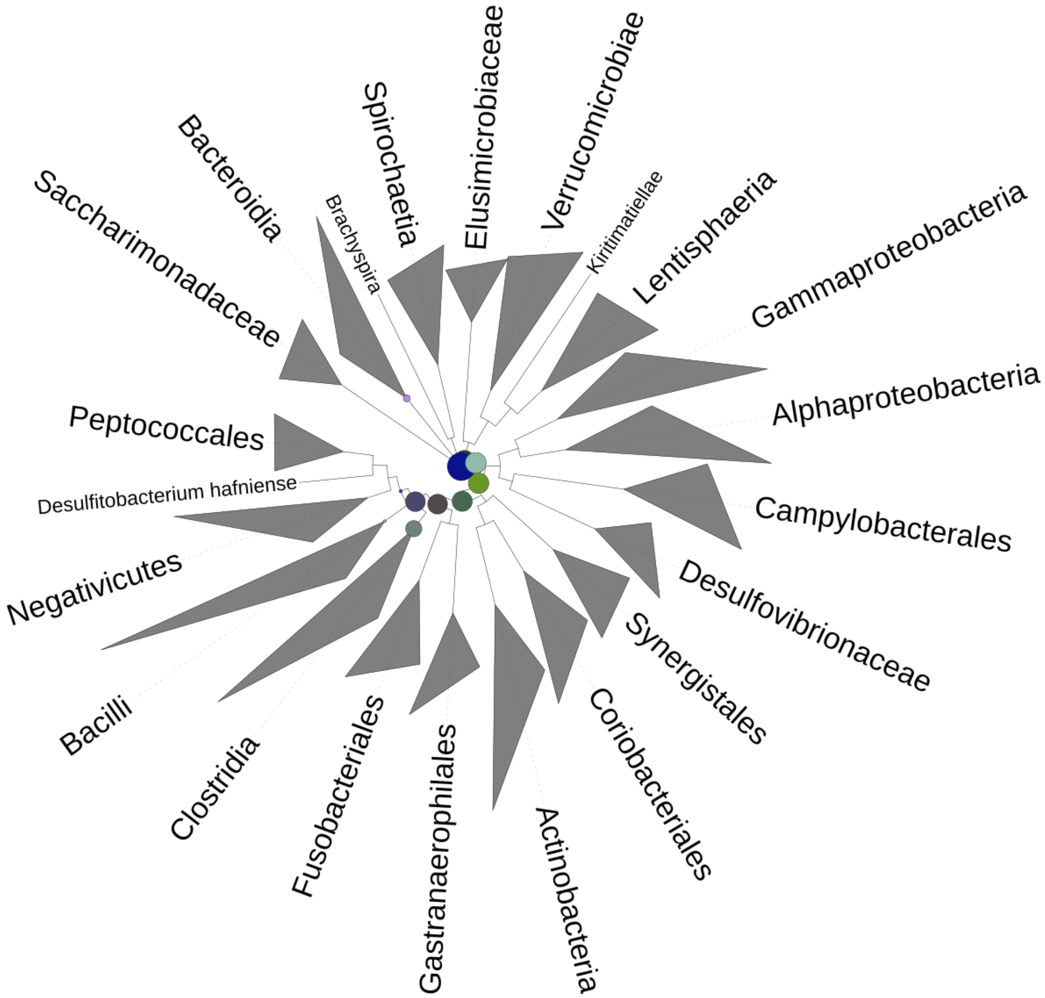


Figure S9: A summary of the species diversity of a metagenome dataset on the gut tree. The circles on the tree represent number of reads mapped to each clade, and their sizes are proportional to the count. Clades are collapsed to class level in this case for clarity.
